# Bacterial glycyl tRNA synthetase offers glimpses of ancestral protein topologies

**DOI:** 10.1101/2021.08.20.456953

**Authors:** Jorge-Uriel Dimas-Torres, Annia Rodríguez-Hernández, Marco Igor Valencia-Sánchez, Eduardo Campos-Chávez, Victoria Godínez-López, Daniel-Eduardo Rodríguez-Chamorro, Morten Grøtli, Cassandra Fleming, Adriana Hernández-González, Marcelino Arciniega, Alfredo Torres-Larios

## Abstract

Aminoacyl tRNA synthetases (aaRSs) are among the proposed proteins present in the Last Universal Common Ancestor (LUCA). There are two types of glycyl tRNA synthetases (GlyRSs), from which the archaeal-eukaryal type is the one suggested to be present in LUCA. Here we solved the crystal structure of a complete bacterial glycyl tRNA synthetase (bacGlyRS) and show that indeed, bacGlyRS carries several structural signals that point it at the origin of all aaRSs. Furthermore, if bacGlyRS is ancestral, it should help to build a reliable Tree of Life (ToL). Given the modular nature of protein evolution, we used only two sub-domain segments with duplicated ancestral topologies, no detected orthologs and an assumed limited horizontal gene transfer (HGT). These motifs correspond to the non-specific RNA binding regions of contemporary bacGlyRS, archaeal CCA-adding enzyme (arch-CCAadd), and eukaryotic rRNA processing enzyme (euk-rRNA). The calculated, rooted bacterial ToL agrees with several phyla relationships unaccounted by the available trees.

## Introduction

Among the 355 genes proposed to have been present in LUCA, eight code for aaRSs, enzymes that attach cognate amino acids to their corresponding tRNAs. Glycyl tRNA synthetase (GlyRS) is found between these candidates (1). Unexpectedly, there are two types of GlyRS that do not share a single common ancestor: an α_2_ homodimer (eukGlyRS), related to HisRS, ThrRS, ProRS, and SerRS and present in eukarya, archaea, and some bacteria and an α_2_β_2_ heterotetramer (bacGlyRS), related to AlaRS and present in bacteria (2), (3), (4). Due to its distribution in the three life domains, it is believed that eukGlyRS is the ancestral type that was present in LUCA (1). However, bacGlyRS is found in bacteria that are deeply rooted, like *Thermotoga* and *Cyanobacteria* (5), (Figure S1). Therefore, the questions regarding why there are two GlyRSs and which one may have appeared first remain unanswered. The structure of bacGlyRS may offer further insights into the molecular evolution of aaRSs, their protein domains, and the origins of the genetic code (6).

Here we show the three-dimensional structure of a functional, complete bacGlyRS from the thermophilic bacteria *Thermanaerotrix daxensis* (Chloroflexi) (7) in complex with glycine. The structure was solved by X-ray crystallography and verified in solution by negative staining, electron microscopy experiments. The structure represents the biggest of all aaRSs, but integrates the smallest known catalytic domains (α, around 30 kDa) and the largest tRNA interacting subunit (β, around 70 kDa). It consists of an α_2_ dimer, with a β-subunit interacting aside each α-subunit. Regarding homology, the β-subunit shows remote relationships with archaeal CCA-adding enzymes-eukaryotic rRNA processing complex, metal-dependent HD phosphohydrolases, and the anticodon binding domain (ACBD) of class I aaRSs, particularly ArgRS and CysRS. All these domains belong to the most ancient groups of enzymes and have sequence identities with bacGlyRS below 10%. Unexpectedly, a model in complex with tRNA shows that the orientation of this substrate would be inverted relative to all class II synthetases, making an hybrid of class I and II type of tRNA recognition. The atypical features found in both the catalytic domain (4) and on the putative complex with tRNA, together with the size of the β-subunit, and the homologies with ancient folds and proteins, demonstrate that bacGlyRS α-subunit represents a primitive aaRSs. The CCA-adding-rRNA processing-like architecture of the β-subunit represents a multifunctional, primordial non-specific RNA binding protein. Together with bacGlyRS, they make an example of the coevolution of the ribonucleoprotein (RNP) world (8). These three proteins are related by a common, duplicated βαββ topology with an ancient ferredoxin-like fold. We calculated a phylogenetic tree using the universe of archaeal and bacterial proteins, most proximal by sequence identity. We thus obtained a rooted tree that agrees with several previous observations. Still, it also suggests novel relationships between species, and it places the most deeply rooted bacteria within the anaerobic, sulfate-reducing *Syntrophobacterales*.

## Results

### - bacGlyRS is a βααβ complex formed around a central α_2_ dimer that is not related to eukGlyRS, forming two different subclasses

To gain further insights into the evolution of bacGlyRS, we solved the crystal structure of the naturally linked *αβ*-subunits of GlyRS of *Thermoanaerotrix daxensis* (see Supplementary Results). The comparison of the two types of GlyRS (bacterial and eukaryal, Figures S5, S6C, D) shows that the organization and relative orientation of the oligomeric ensembles is quite different. In eukGlyRS and most class II aaRSs, besides the catalytic domain, other domains of one monomer are involved in interface contacts to stabilize the oligomer and/or make intersubunit contacts with the tRNA (Figure S6D). In contrast, bacGlyRS represents a unique homodimeric dependence on merely the catalytic domain, where each β, tRNA-interacting subunit, is completely independent of the homodimer (Figures 1, S5 B, D). The bacGlyRS α_2_β_2_ heterotetramer would be better defined as a βααβcomplex formed around a central α_2_ dimer (9). In addition, none of the domains shares structural homology between the GlyRSs, except for the core of the catalytic domain, with an extremely limited similarity between the two types of GlyRS. It has been shown that even in this domain, eukGlyRS is more similar to subclass IIa enzymes than to bacGlyRS (Valencia-Sánchez et al., 2016). Different subclasses must define the difference between the two types of GlyRS. Somewhat surprisingly, none of the five domains of the β-subunit is clearly shared between the subclass IId members, bacGlyRS and AlaRS, restraining the homology between these enzymes to the features shared in the catalytic domain (4).

**Figure 1.**
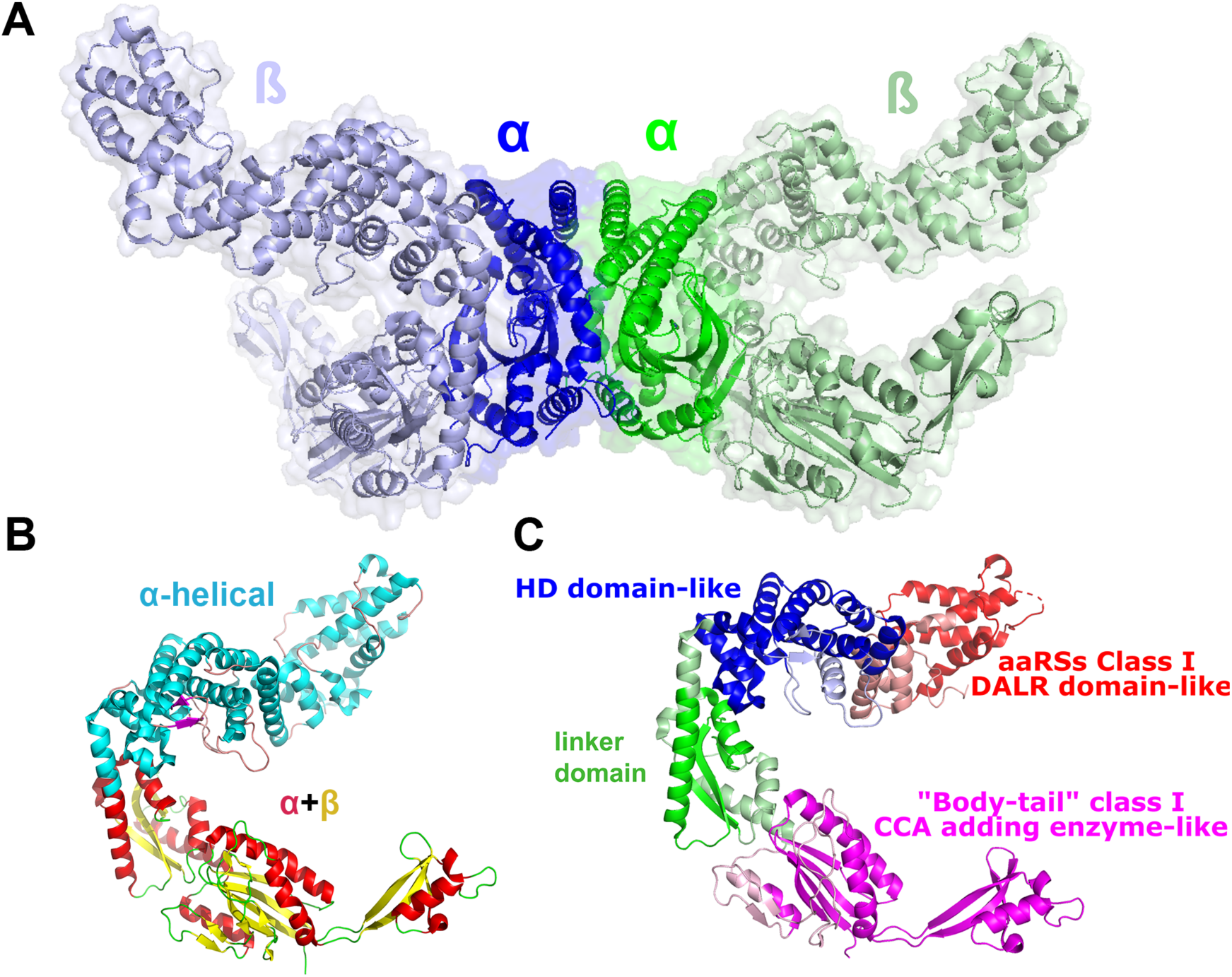
bacGlyRS is a βααβcomplex formed around a central α_2_ dimer. The β-subunit possesses two types of architecture, each with domains homologous to tRNA binding proteins specific for the acceptor (ancient) and anticodon stems. **A.** Each α subunit (300 aa) is flanked by one β subunit (700 aa), making bacGlyRS the biggest known aaRS of all three life domains (see Supplementary Discussion). Remarkably, the α-subunit represents the smallest known class II catalytic domain, and thus the β-subunit, which is easily dissociated *in vitro*, provides most of the molecular size of the enzyme. **B**. The two architectures of the bacGlyRS β-subunit. In cyan is depicted an almost all α-helical region that defines the HD phosphorylase-like domain and the tRNA anticodon binding domain of class I aaRSs (ACBD or DALR). These domains are shown in blue and red in **C,** respectively. In yellow and red is an α+*b* region that comprises three domains: a linker domain (green in **C**) with no clear homology to any protein, although the closest structural match is with the palm domain of DNA polymerases, and two domains with clear homology to class I, tRNA CCA adding enzymes from archaea (pink in **C**). **C**. The five identified domains in the bacGlyRS β-subunit are shown in different colors.

### - The homology of bacGlyRS β subunit with tRNA binding domains allows to confidently build a model of the aaRS-tRNA complex

To gain further insight into the evolutionary origin of bacterial GlyRS, we performed structural searches using the algorithm Dali and the DASH database. We gathered the most representative models showing homology for each domain, trimmed the corresponding regions, aligned them using the STAMP algorithm, and clustered with Bio3D and BioVinci. We also performed Principal Component Analysis (PCA) on the Cα atoms of the core residues found, according to the algorithm implemented in Bio3D. Surprisingly, the only clear structural homologs that we found (Figures 1C, 2A, S8-S9, Table S3) are the class I anticodon binding-domain (ACBD or DALR domain, in particular with ArgRS and CysRS, Figure S8), phosphohydrolases possessing the HD domain (Figure S9) and the body-tail domains of archaeal tRNA CCA adding enzymes-eukaryotic rRNA processing enzymes (Figure 2A, PDB codes 1R89, 1SZ1 (10) (11) and 4M5D (12), respectively).

**Figure 2.**
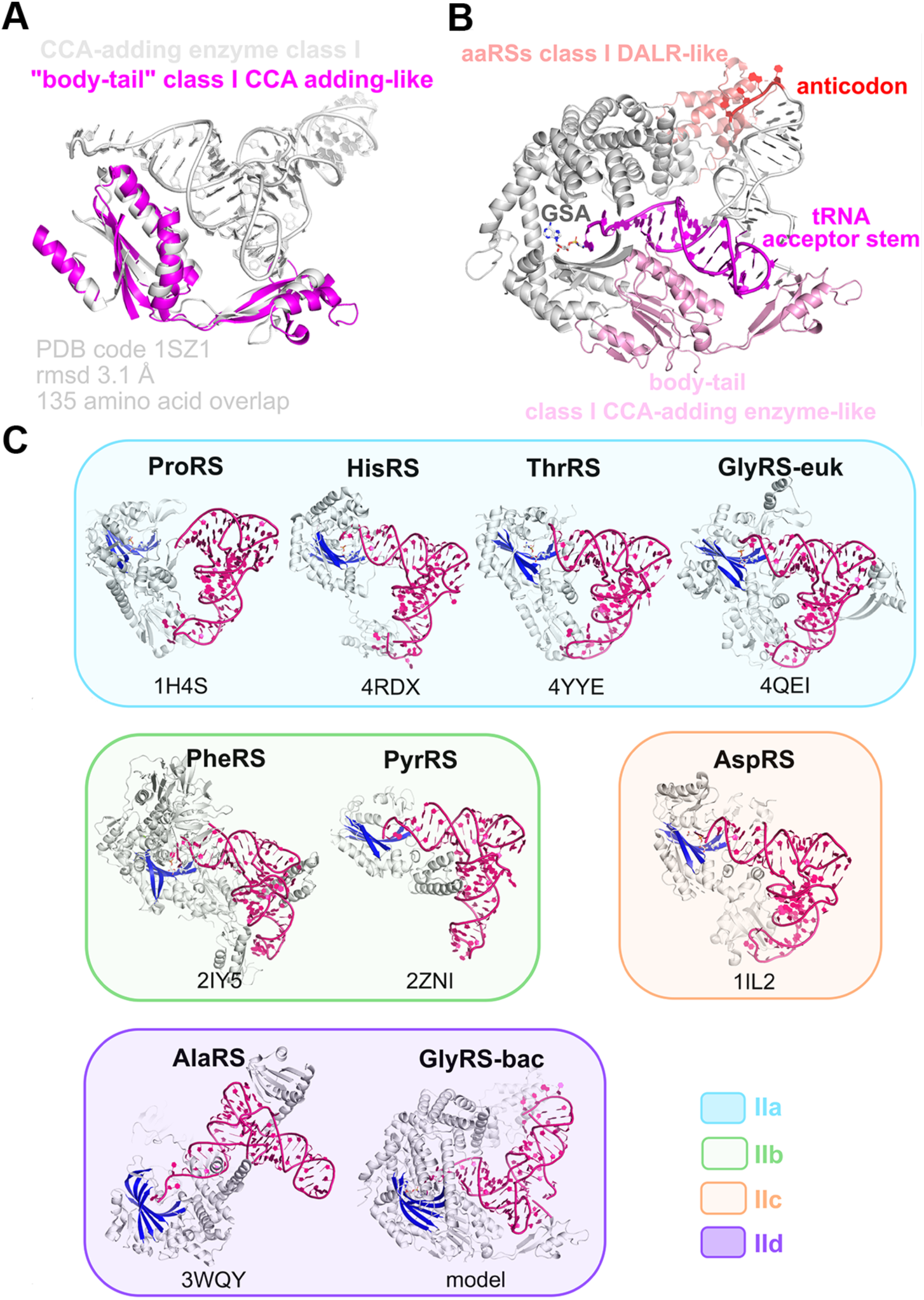
The archaeal CCA-adding enzyme provides a template to build a clash-free model of the bacGlyRS:tRNA:GSA ternary complex. The complex shows that bacGlyRS and AlaRS clearly define a subclass among class II aaRS. **A.** Superposition of the body-tail domains of the CCA-adding enzyme:tRNA complex (white) on the equivalent domains of bacGlyRS (magenta). **B.** Resulting model of the superposition at left but depicted on the whole monomer of bacGlyRS and using the α-subunit model (PDB code 5F5W) to depict the GSA (glycyl-sulfamoyl-adenylate) molecule on the catalytic cavity. The tRNA model is within 6 Å reach of all three regions (body-tail, ACBD, and GSA) and thus fits perfectly on bacGlyRS. **C.** Besides their remarkable similarities at the catalytic domain (Valencia-Sánchez, 2016), bacGlyRS and AlaRS (magenta, subclass IId) would recognize their cognate tRNA in a completely different orientation regarding all other class II aaRSs (see also Figures S6C, D, S7). The relative orientation of the catalytic domain (depicted in part in blue) is the same for all models. The PDB entry code is shown below each model.

The homologies found for the β-subunit allowed us to propose a bacGlyRS:tRNA model (Figure 2B), mostly based on the complex of archaeal CCA-adding enzyme with tRNA (Figure 2A). The superposition of the PDB code 1SZ1 with the coordinates of bacGlyRS (PDB code 7LU4) (Figure 2A) renders a putative complex that interacts with the ACBD and leaves the terminal A76 of tRNA within reach to the glycine moiety of the catalytic subunit (Figure 2B, Figure S6C). Perhaps the most outstanding features of this model are its natural fitting, as given by just placing the template model and, most of all, the resulting relative orientation of the tRNA. Remarkably, if correct, the tRNA recognition would be a hybrid between class I-like type of recognition on a class II aaRS (Figure S7). According to their tRNA recognition, atypical for class II aaRS, bacGlyRS and AlaRS could represent the most ancient subclass of aaRSs, as they may illustrate primitive, somewhat alternate recognition modes given before the class II aaRS:tRNA canonical interaction (Figure 2C). This model further strengthens that bacGlyRS and AlaRS clearly define a subgroup of aaRSs themselves (Figure 2C).

### - bacGlyRS β-subunit presents two architectures: a “recent” α-helical and a putative primitive α+*b* that may be the result of a triplication event

From the structural analysis of the β-subunit, we defined two different architectures (Figure 1B): an α-helical, comprised of domains related to the anticodon recognition and with clear sequence signatures from the HD and ACBD domains, making this region probably more recent than the second seen architecture (Figure 1B, Table S2), which is an α+*b* type, and that only shows structural homology with RNA recognition molecules (Figures 1C, 3D). The finding of a βαββ topology on the three domains comprising the α+*b* architecture (Figure 3A, B) hints that a triplication event was at the origin of this multidomain protein very early in evolution. This hypothesis makes sense because the ferredoxin-like fold is proposed to be very ancient (13), (14), (15), (16) as well as the tRNA acceptor stem (17), (18). Interestingly, most ribosomal proteins from the large subunit are of the α+β type, with ferredoxin-like βαββ topology, also like the RRM fold, typified by the splicing protein U1A (19)

**Figure 3.**
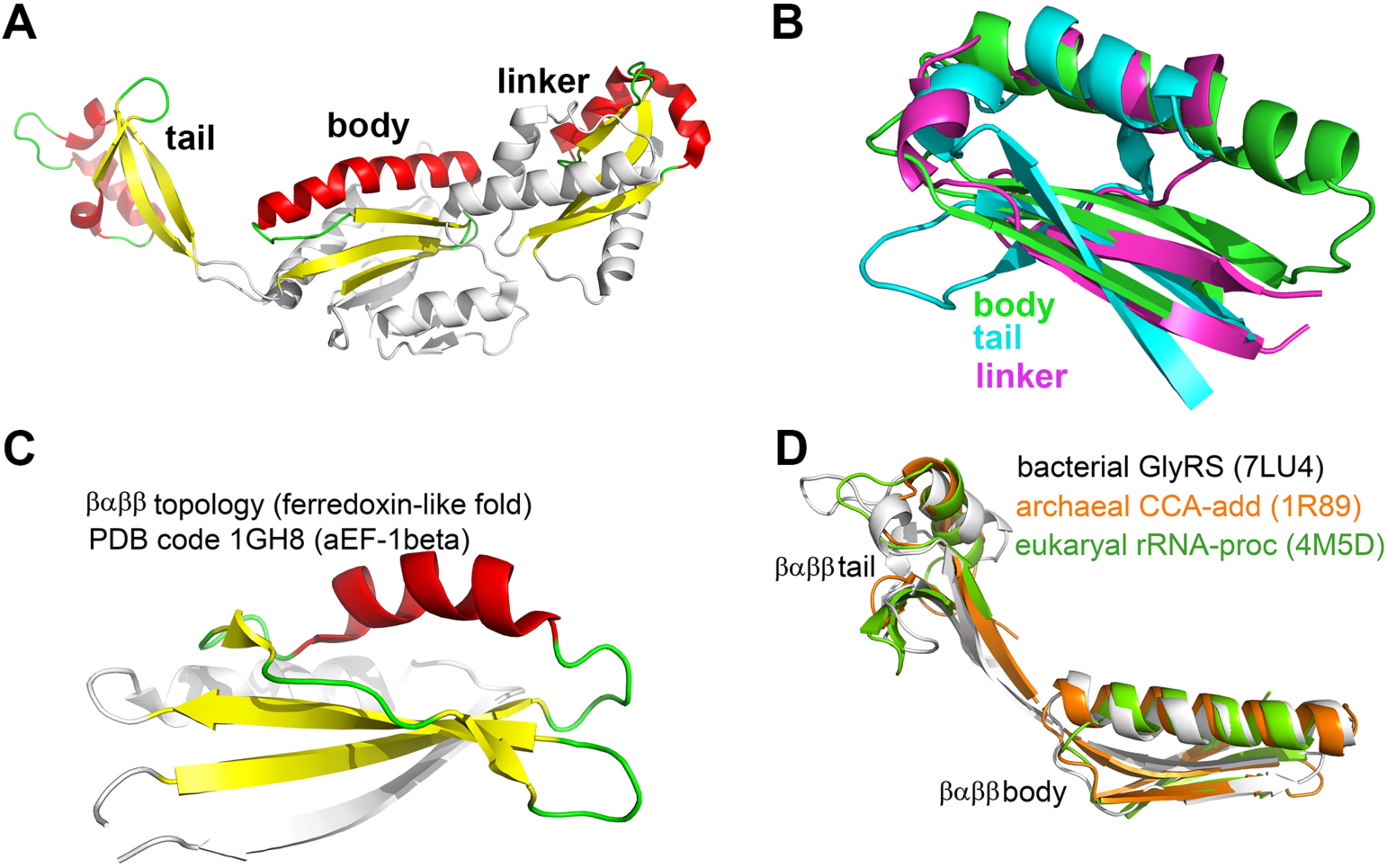
The body-tail-linker domains of the bacGlyRS β-subunit may be the result of a triplication event. The duplicated βαββ, ferredoxin-like, primitive RNA binding topology in the three life domains provides a clear coevolution signal of an ancient RNP world. **A.** The linker, body, and tail domains of bacGlyRS β-subunit show a repeated βαββ topology that can be roughly superposed, as shown in **B.** This topology is homolog of the ancient and widespread ferredoxin fold, exemplified in **C** by the PDB code 1GH8, representing an archaeal elongation factor. This topology is the closest homology of the linker domain with any known protein. That is, it is more similar to other segments of the same chain than to other proteins. Regarding the body domain, it is remarkable that the protein chain traces the βαββ topologies of the body and tail domains and then forms the rest of the body domain with large parts of secondary structures (Figure S10) (43). We also may refer to the body-tail topology as a protein tandem repeat of class V (44). **D.** Superposition of the duplicated βαββ topologies of bacGlyRS, archCCAadd and eukrRNAproc, all representing ancient protein topologies binding to ancient RNA motifs from the three domains of life. The homology between bacGlyRS and archCCAadd can be marginally detected by sequence similarity, around 10%. No other tandem topology homolog has been found in nature, which shows a limited horizontal gene transfer (HGT). This duplicated, not single, topology, reassures its uniqueness and highlights its phylogenetic signal.

### - The duplicated βαββ, ferredoxin-like, primitive RNA binding topology present in the three life domains provides a clear coevolution signal of an ancient RNP world

The duplicated βαββ topology found represents a promiscuous entity but with a single primitive, non-sequence-specific RNA binding activity when found in tandem. This activity was frozen in time because although it is found in the three domains of life, it has been used for single different purposes regarding the binding of ancient RNAs (tRNA acceptor stem and/or rRNA). Thus, we propose that the RNA binding, βαββ tandem topology is an example of the origin of proteins themselves and that their presence in a wide range of different proteins, but with similar ancestral activity, was raised by a single, extremely ancient HGT event, placing the origin of the three domains of life in a very tight window of time. The tandem topology itself may be of pre-LUCA (Last Universal Common Ancestor) origin.

### - The ancestral, duplicated βαββ, ferredoxin-like topologies of archaea-CCAadding and bacterial β-GlyRS draw the rooted bacterial Tree of Life

With the hypothesis described above, we calculated a phylogenetic tree (Figure 4) using the sequences of the available βαββ duplicated topologies of archaea and bacteria domains. Notably, we could use the Archaeal, CCA-adding enzyme sequences to obtain a rooted tree (Figure 4) that yields information consistent with the widely accepted taxonomy, like the overall, Gracilicutes vs. Terrabacteria division (20), the general Firmicutes phylogeny division of Limnochordales, Halanaerobiales, Negativicutes (21), or the placement of Magnetococcales as an Alphaproteobacteria outgroup (22). It also proposes that Methanopyrus is the closest Archaea related to bacteria (23); to divide Deltaproteobacteria into four phyla that do not belong to proteobacteria (24). Epsiloproteobacteria is also suggested to be reclassified as not a Proteobacteria, as proposed in (25). Notably, our proposal implicates that Thermotogales should be considered members of the Firmicutes, as already suggested by (26) and places Fusobacteria very close to the root of the tree, as recently proposed (20).

**Figure 4.**
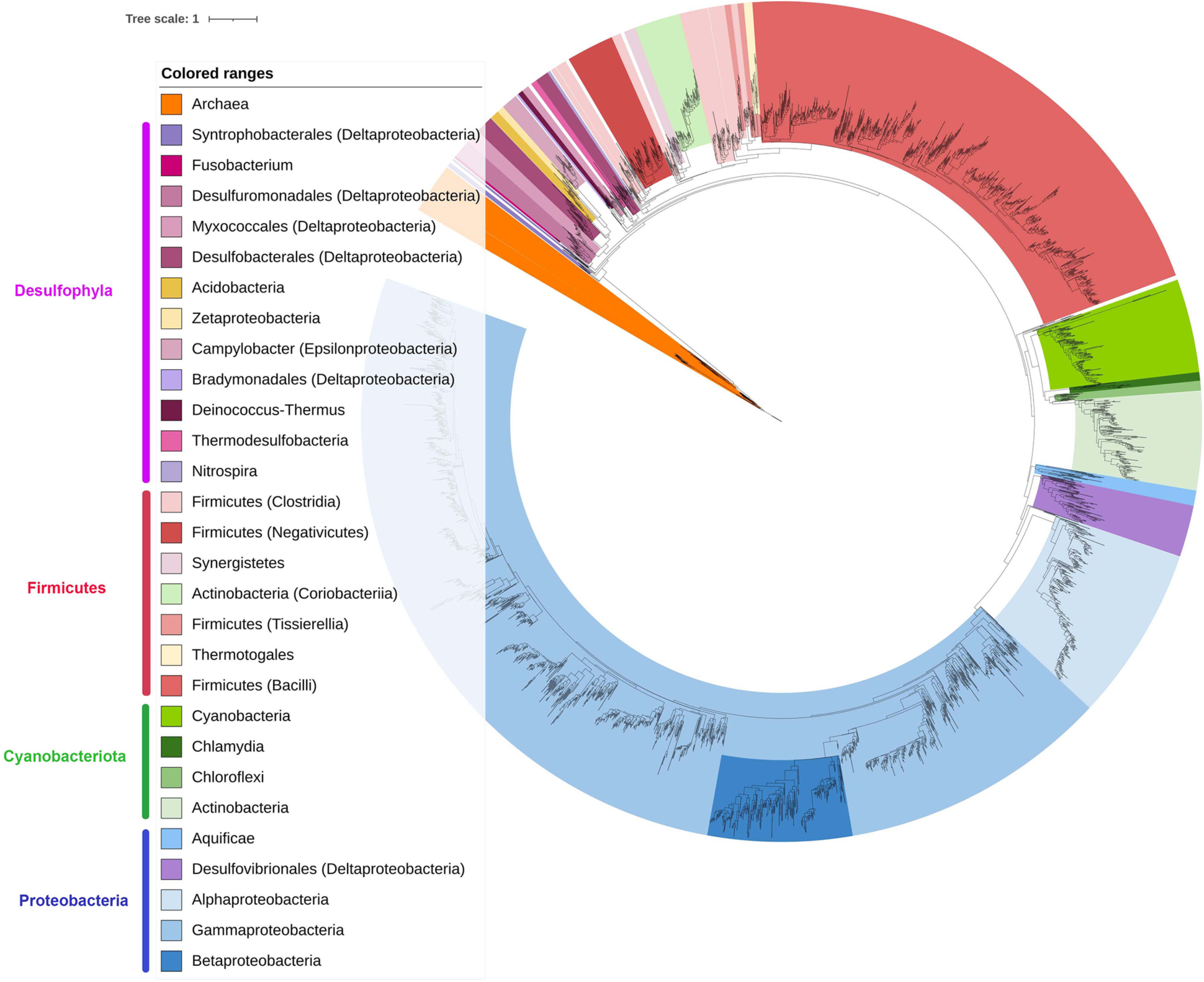
The ancestral, duplicated βαββ, ferredoxin-like topologies of archaea-CCAadding and bacterial β-GlyRS draw the rooted bacterial Tree of Life. The expected limited Horizontal Gene Transfer (HGT) of the duplicated βαββ topology of bacGlyRS led to this proposition, rooted in the archaeal βαββ topology of archaeal CCA-adding enzyme. Several deeply rooted phyla belong to what was previously known as Deltaproteobacteria. We propose four superphyla, with Desulfophyla as the root. Fusobacterium, recently proposed to be close to the root, remains within this group. The tree was calculated with IQ-TREE v.2.1.3. (45), with no sequence outliers in the composition test. The diagram was drawn with iTOL. The tree and the sequences from which it was derived are available in the Supplementary Material.

Perhaps our proposed tree’s two most outstanding features are i) the relationship between Cyanobacteria, Chlamydia, Chloroflexi, and Actinobacteria. Interestingly, Cyanobacteria and Chloroflexi are two of the seven phyla where reaction-center phototrophy has been reported (27), (28), (29). Also, the relationship between Cyanobacteria and Chlamydia has been reported (30), (31), (32), although also disputed (33). Finally, Actinobacteria seems to be somewhat related to photosynthesis, as already suggested (34). Remarkably, no other available phylogenetic tree opens the possibility that these four phyla are related; ii) Deltaproteobacteria (sulfur and sulfate-reducing bacteria) are placed in the root of the bacterial ToL. It has been proposed that the deepest branches of the phylogenetic tree comprise chemolithotrophic autotrophs. The first autotrophic processes were akin to those seen in the sulfur and sulfate-reducing bacteria (35).

## Discussion

GlyRS is supposed to be one of the first aaRSs to evolve. But what motivated the appearance of two GlyRSs? The structure of bacGlyRS offers a possible explanation. The α-subunit of bacGlyRS may have appeared as one of the first aaRSs (Table S2), even in a pre-LUCA world, where the *αβ* architecture of the β-subunit served as a multifunctional, helical-RNA binding/recognition module. Perhaps this recognition was based on size only, not sequence, as it is for the CCA-adding enzyme. As the genetic code and the tRNA became more sophisticated, other domains, including the ACBD, were created or recruited in bacteria. At this time another, full GlyRS, smaller and including both the catalytic and the anticodon recognition regions (eukGlyRS), rapidly evolved in archaea from ProRS, or another class IIa aaRS, that substituted the primordial, α-bacGlyRS and was inherited, post-LUCA, to some bacterial species through horizontal gene transfer (HGT). It is possible that archaea inherited from LUCA the primordial bacGlyRS to quickly lose it afterward, leaving some domains of the β-subunit (i.e., the tail and body) useful for other enzymes (i.e., the CCA-adding).

Given the relationships of bacGlyRS with AlaRS (catalytic core and primordial aaRS) and ArgRS (primordial ACBD), it is possible to map them on the genetic code table (Figure 5) and obtain a correlation between the emergence of these putative primordial aaRSs with the tRNA acceptor stem code (first code, second codon letter) and the anticodon stem code (second code, first codon letter). Remarkably, it is not possible to make such a relationship with any other aaRS, which further highlights the ancient nature of bacGlyRS. We chose to perform our phylogenetic analysis using the duplicated βαββ topology found because it may represent an example of ancient domain accretion and multiple domain repeats not identified by sequence. Ancient proteins described tend to be a single-domain, so we consider that the duplicated, sub-domain topology offers a better chance of unveiling truly ancient evolutionary events.

**Figure 5.**
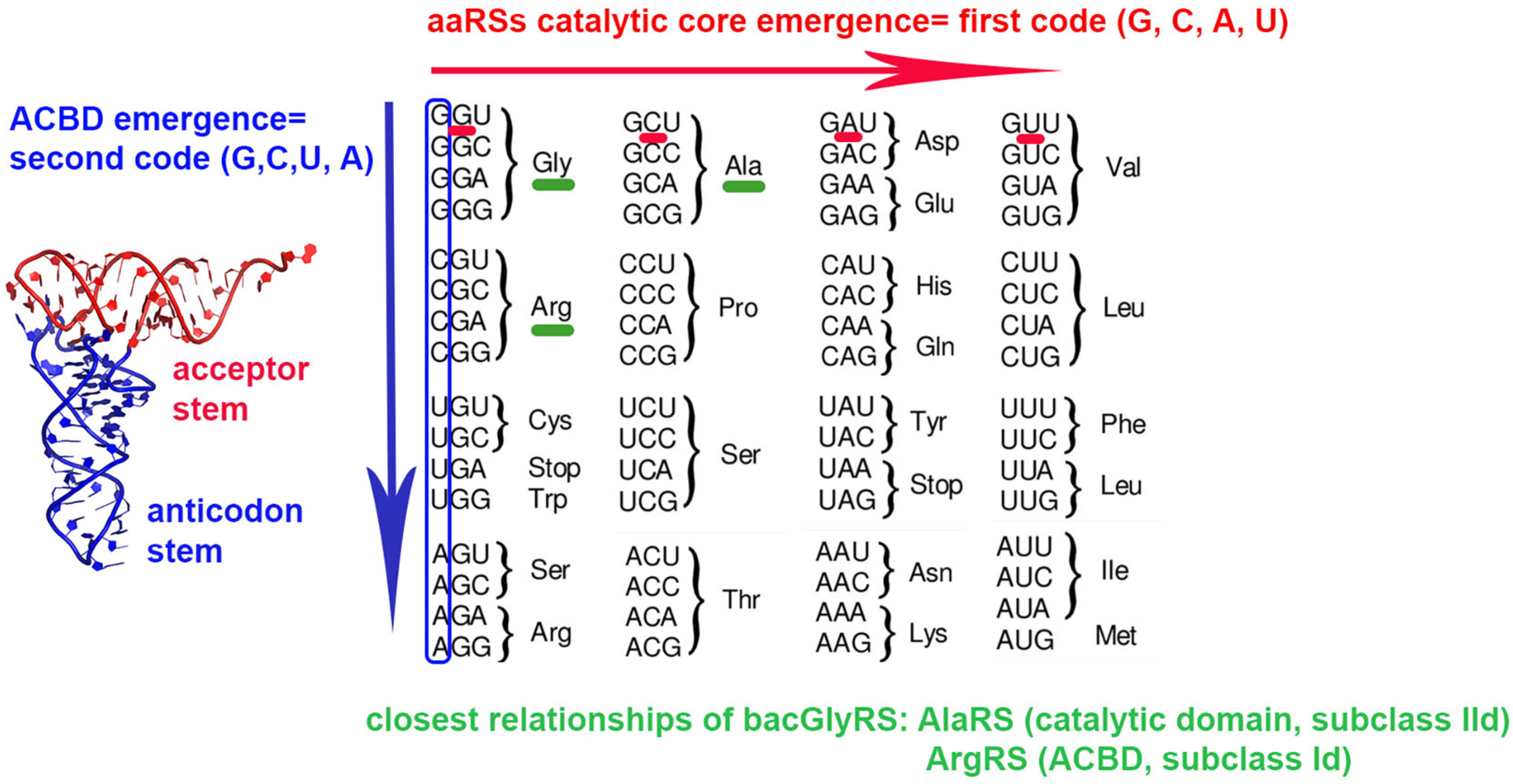
The genetic code table reflects the age of primordial aaRSs and their evolutionary reading (second base first, first base second). The “first, or operational code” (46) corresponds to the second letter of the codon (47), and it is suggested to lie on the acceptor stem of the tRNA (ancient). A complex between an aaRS catalytic core and the βαββ body-tail domains may have carried out the primitive aminoacylation task. The “second code” corresponds to the first letter of the codon and would lie on the anticodon stem of the tRNA. This proposal agrees with an almost concomitant appearance of class I/II aaRSs and previous hypotheses on the emergence of aaRSs (48). The first, primitive code, could have been specific for Gly, Ala, Arg, and Pro.

Overall, this work proposes (see Highlights in the Supplementary Material) that a clear phylogenetic signal may be extracted from single, carefully chosen ancient protein fragments. It illustrates the coevolution of a ribonucleoprotein world, in which protein ancestral topologies were duplicated to generate multidomain proteins able to bind promiscuous RNAs. It also represents a step towards deciphering the grammar of primordial domain combinations and how the very first proteins emerged.

## Experimental procedures

Refer also to Supporting information

## Crystallization and structure determination

The crystallization test was performed by placing 1.5 μL of protein [36 mg/mL], 1 uL of the corresponding dilution, and 1.5 uL of mother liquor (550 mM magnesium formate, 1% dextran sodium sulfate M_r_ 4,000, 100 mM glycine pH 8.5, 3% sucrose, 100 mM sodium acetate pH 4.6). Diffraction data were collected at 100 K using a wavelength of 0.9785 Å at the Life Sciences Collaborative Access Team (LS-CAT) 21-ID-D beamline at the Advanced Photon Source (Argonne National Laboratory, Argonne, Illinois, USA). Data were indexed and processed with xia2 (https://xia2.github.io/) that used XDS (36), Pointless (37) and reduced with Aimless (38). The structure was solved by molecular replacement using MrBUMP (39), with a search model based on PDB entry 5F5W. The initial model was obtained using the Phenix.Autobuild program of the PHENIX package (40) and was displayed and manually built in COOT (41). Refinement was alternated with the use of the PDB_REDO server (42).

## Data availability

The atomic coordinates and structure factors of the *αβ* bacGlyRS from *T. daxensis* have been deposited in the Protein Data Bank (PDB), code 7LU4.

## Supporting information

sequences-for-bacterial-tree

bacterial-tree

bacterial-tree-colors

bacterial-tree-IQtree-log

putative-complex-tRNA-bacGlyRS

pdb-aligned-archaea

pdb-aligned-eukarya

pdb-aligned-bacteria

dimeric-GlyRS-to-map-with-colors

dimeric-GlyRS-colors

## Supporting information

This article contains supporting information.

## Acknowledgments

The authors thank Zdzislaw Wawrzak for help with the crystallographic aspect of this project, Elizabeth Nallely Cabrera González, Georgina E. Espinosa Pérez, Pamela J. Focia and Valerie Tokars for technical guidance.

## Author contributions

A.T.L, A.R.H., M.I.V.S, and D.E.R.CH conceived the project. J.U.D.T and V.G.L. purified and crystallized TdGlyRS. A.R.H performed the activity experiments. A.T.L. collected crystallographic data and solved the structure of TdGlyRS. E.C.CH. performed and analyzed the EM negative staining experiments. M.G. and C.F. synthesized the GSA compound. E.C.CH. and V.G.L. performed the affinity measurements. M.A., D.E.R.CH., A.H.G., and A.T.L. performed the refinement of the model and the structural and sequence analysis. A.T.L. and A.R.H. wrote the manuscript.

## Funding and additional information

A.T.L. was supported by Universidad Nacional Autónoma de México (PAPIIT-DGAPA) grants IN202416 and IN204820. Use of the Advanced Photon Source was supported by the U.S. DOE under Contract No. DE-AC02-06CH11357. Use of the LS-CAT Sector 21 was supported by the Michigan Economic Development Corporation and the Michigan Technology Tri-Corridor, Grant 085P1000817.

## Supplementary material

### Highlights

Working hypotheses

1. There may be a link between proteins dealing with the genetic code (present in LUCA), tRNA, and the appearance of the first proteins (1), (2).
2. The first multidomain proteins may have evolved from duplication events (3), (4), starting from peptides and sub-domain motifs (5), (6), (7), (8).
3. As gene transfer is mostly vertical in bacteria (9), it may be sufficient to choose a single, actual ancestral protein fragment, to uncover the evolution of this domain of life.

Facts provided by this work

1. We solved the crystal structure of a putatively ancestral protein, bacGlyRS (Table S2).
2. bacGlyRS is unique to bacteria and is the only aaRS that delineates its two functions because it possesses two subunits that separate the catalytic domain (α) from the tRNA recognition regions (β) (Figure 1A).
3. The β-subunit has two structural regions (α and α+*b*) that correlate by age with the two tRNA recognition regions (Figure 1B, C). These regions may be a clear example of the coevolution of the RNP world (Figure 3).
4. The α+*b* region has two βαββ primitive topologies in tandem found in the three domains of life, in different proteins, but performing the same RNA, non-sequence specific, binding function (Figure 3).

Novel hypotheses

1. The genetic code table reflects the age of primordial aaRSs and their evolutionary reading (second base first, first base second) (Figure 5).
2. Bacterial species that possess bacGlyRS should be more ancient (Figure 2, Discussion and Supplementary Discussion).
3. The βαββ tandem topology is the result of a pre-LUCA duplication event aimed to coevolve with the RNA world.
4. If bacGlyRS is ancient, it is possible to build a trustful phylogenetic tree (extremely limited HGT -only present in the bacterial domain, with no paralogs-) using the most ancient part of the protein.
5. We used the βαββ tandem topology to build a tree, which is possible to root using the archaeal, CCA-adding enzyme ortholog (Figure 4).

## Supplementary Figures

**Figure S1.**
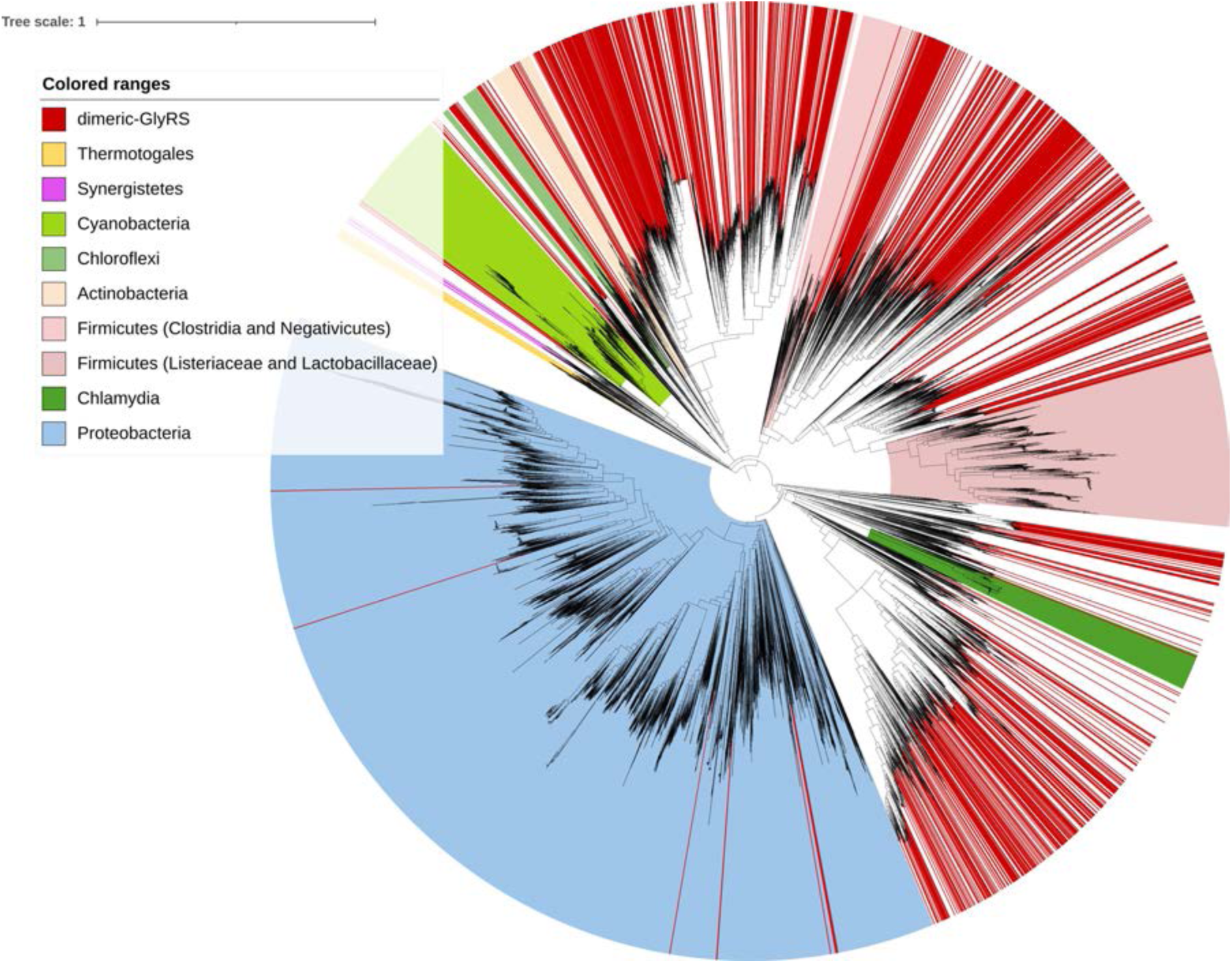
bacGlyRS is exclusively present on particular bacterial clades and superphylum. In red are the bacterial organisms with the dimeric GlyRS. In other colors are the clades that only (or mainly) present the bacterial GlyRS. Bacteria that possess the dimeric, eukaryotic-type GlyRS, are marked in red and taken from EggNOG Database COG0423 http://eggnog5.embl.de/#/app/home. The diagram was adapted from the data published in (10) and drawn with iTOL (11). Candidate Phyla Radiation (CPR) (12) seems to possess mainly the dimeric GlyRS. These phyla would be more recent than bacteria with bacGlyRS, according to our proposed model.

**Figure S2.**
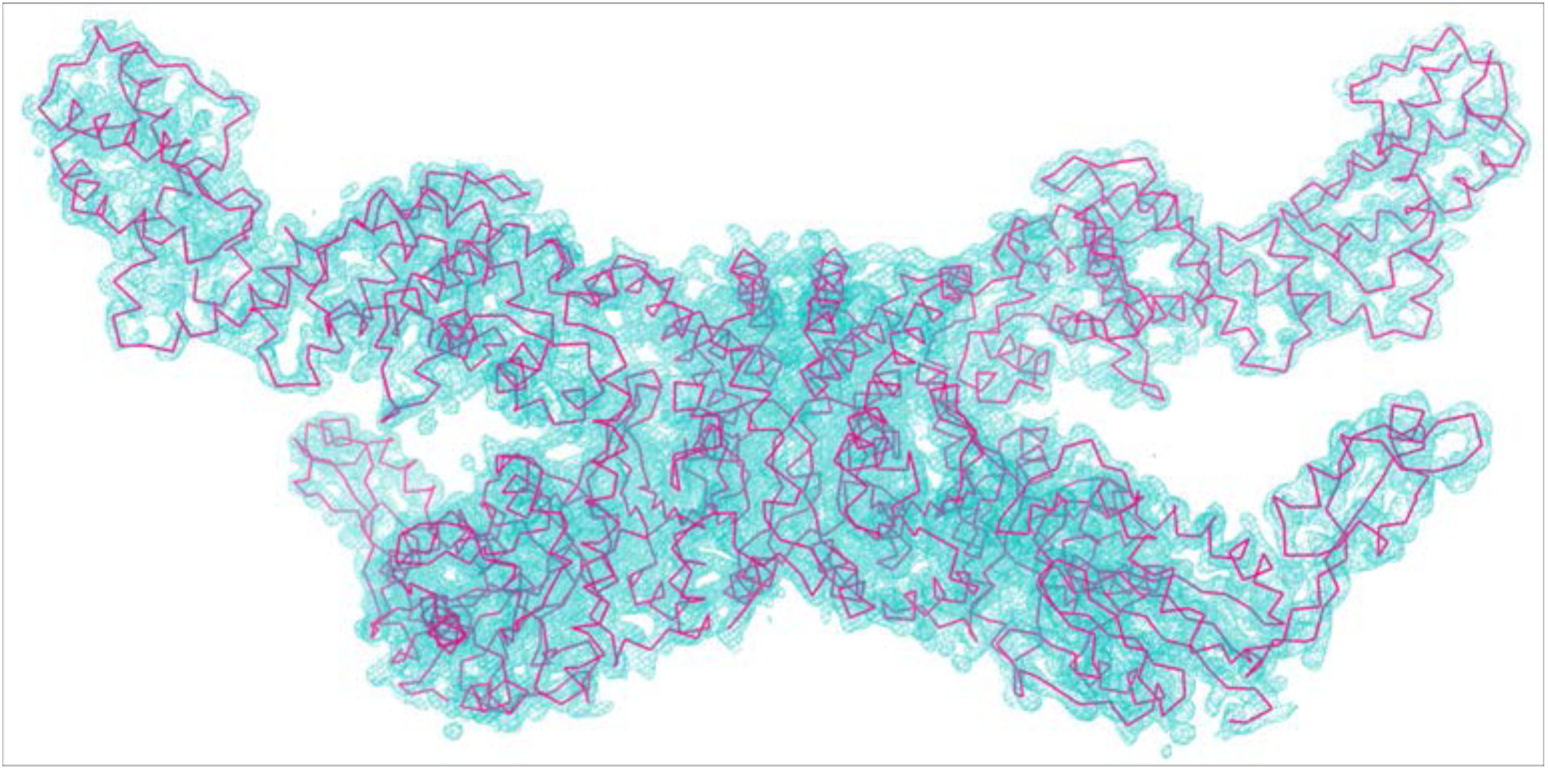
The experimental electron density allows modeling the complete, 2000 amino acid bacGlyRS model. Simulated annealing, composite omit map (cyan) (13) contoured at 1.5 *σ*. The built model, PDB code 7LU4, is depicted in red.

**Figure S3.**
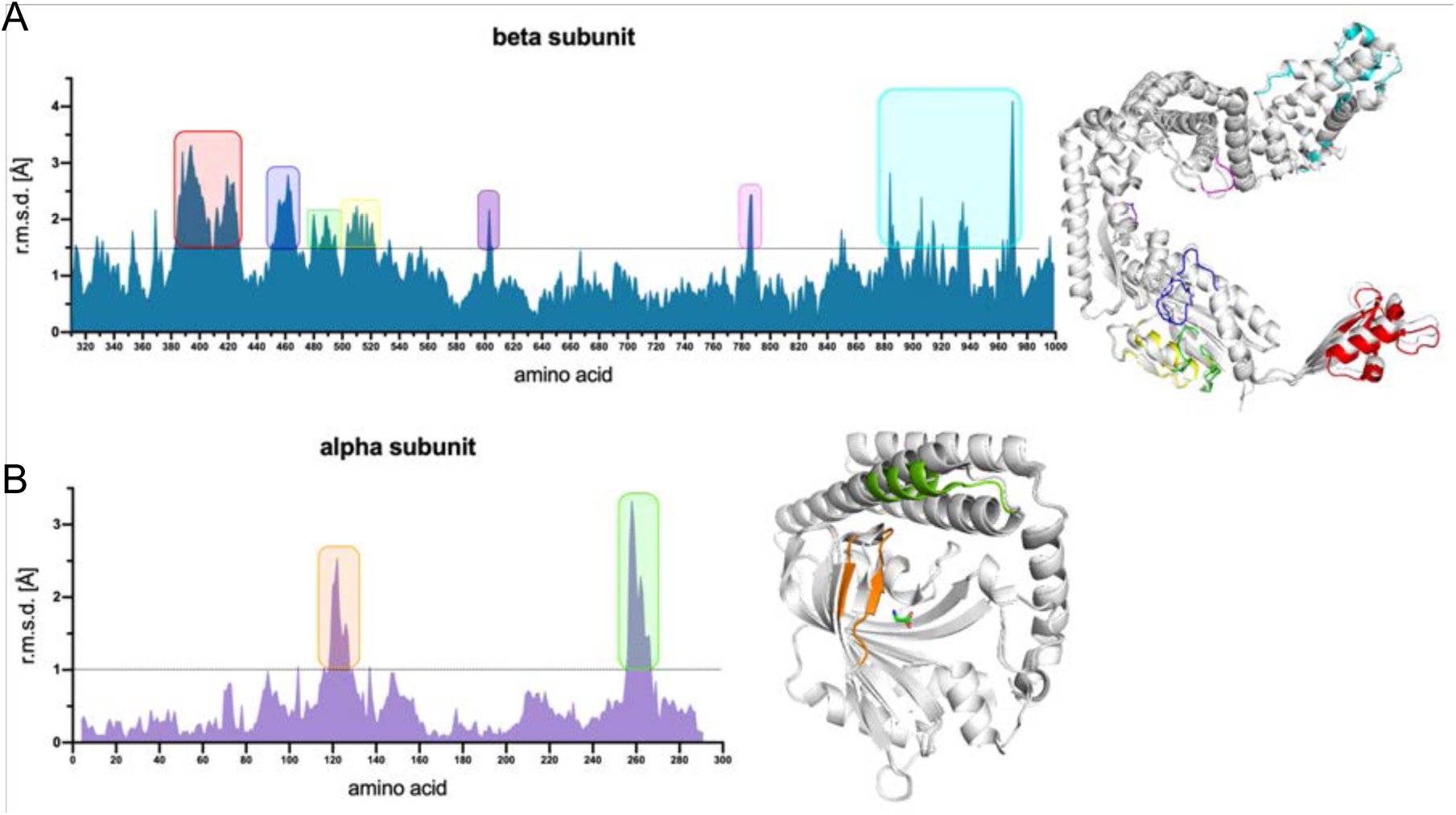
The binding of glycine on bacGlyRS promotes the same conformational change as the binding of GSA on the α-subunit (PDB code 5F5W). The asymmetric unit of PDB code 7LU4 reported in this work contains a dimer with one monomer in the presence of glycine. The Figure depicts the superposition of the *αβ* apo monomer on the *αβ* monomer in complex with glycine. **A**. The β subunit presents some movements in the tail (red) and ACBD domains (cyan). **B**. The region in orange (present both in bacGlyRS and AlaRS) contains a conserved Trp in the α- subunit that interacts with the amino group of glycine through a cation-*π* interaction (14).

**Figure S4.**
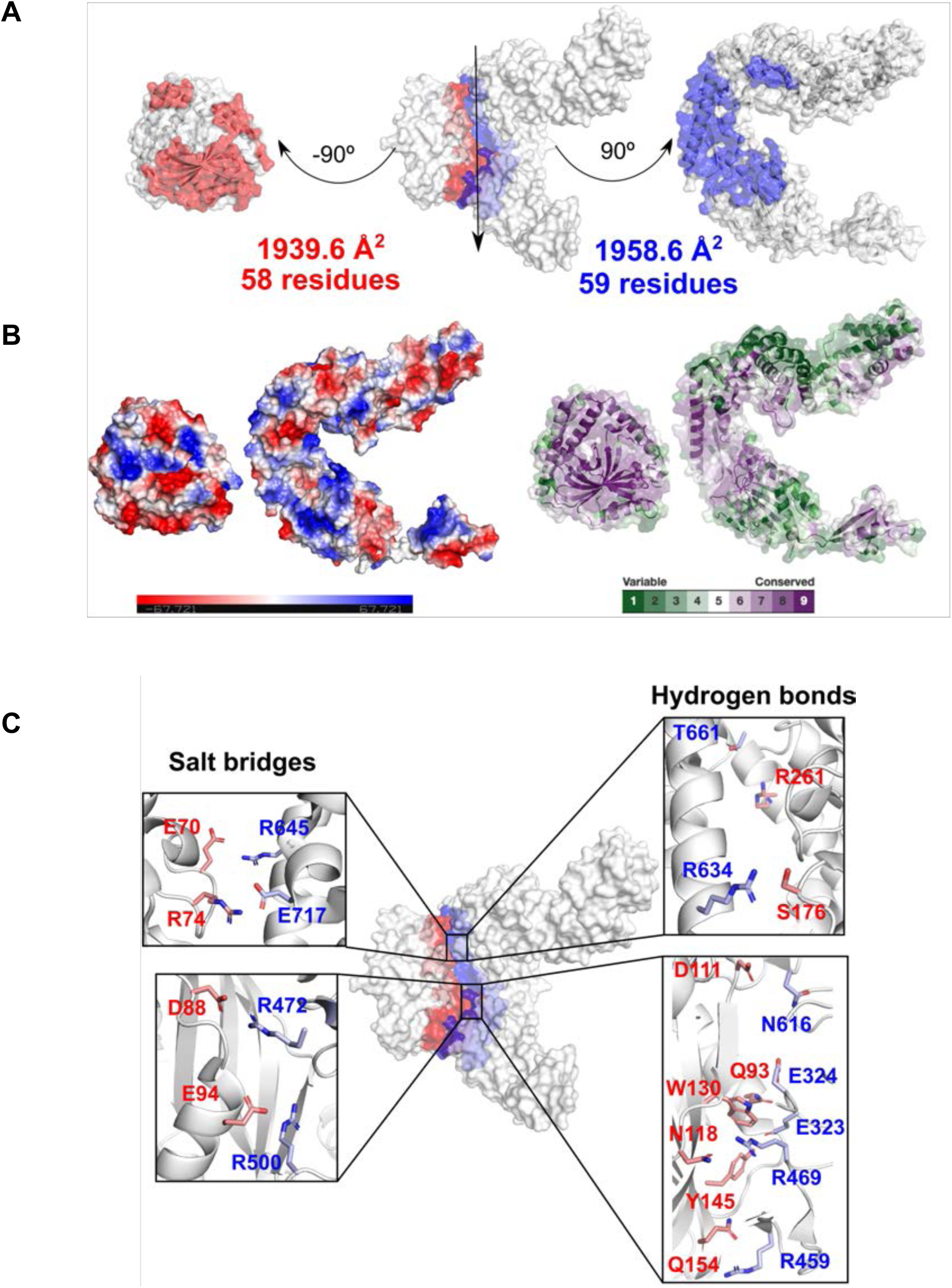
The *α-b* subunit interactions are highly complementary. **A.** The *αβ* interacting area covers almost 2000 A^2^. **B**. The electrostatic surface and the conserved residues are in excellent agreement, and **C**. Several salt bridges and hydrogen bonds stabilize the non-covalent complex. These interactions validate the biological relevance of the observed relative orientation of the *α-b* subunits.

**Figure S5.**
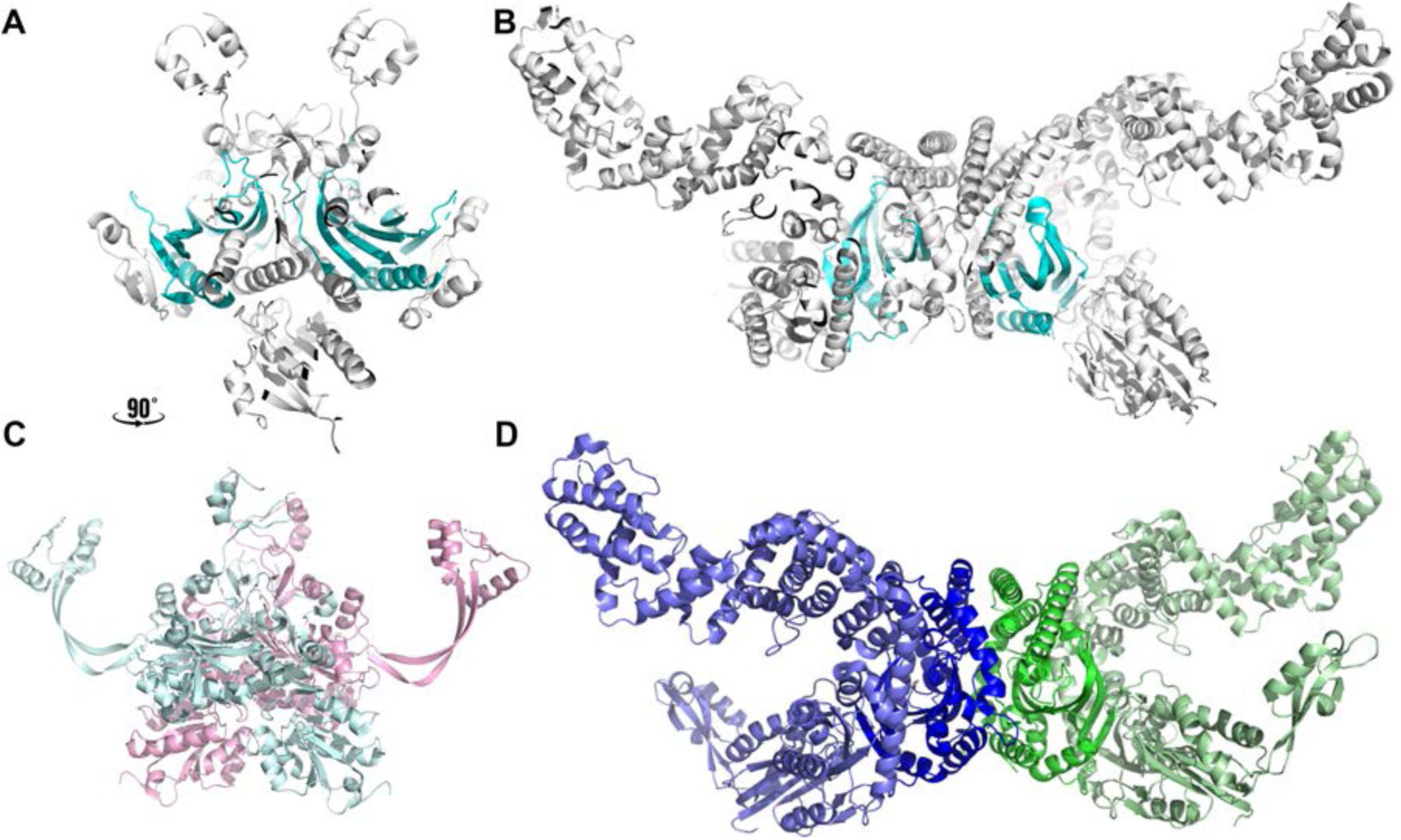
The two types of GlyRS are not related, and therefore they belong to different subclasses. They only share the general fold of the catalytic domain (cyan). **A**, **C** Two different views of eukGlyRS (PDB entry 4QEI). **B**, **D** bacGlyRS. A and B show the same orientation of the catalytic domain (cyan). bacGlyRS has the shortest catalytic domain among all aaRSs. **C** and **D** show a better view of the domains where tRNA would bind (see below). Remarkably, the orientation of bacGlyRS remains almost unchanged (see below).

**Figure S6.**
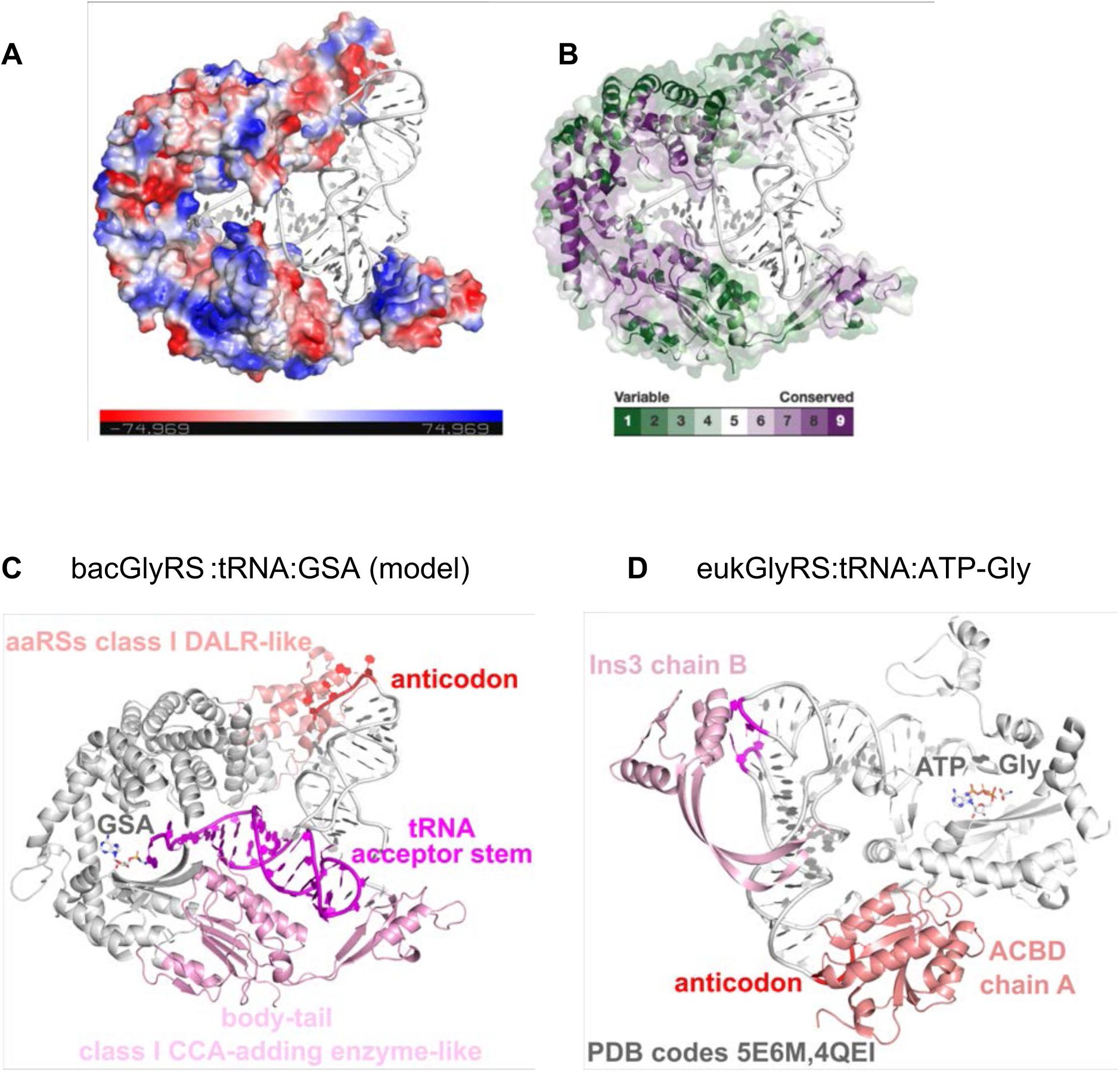
The two types of GlyRS are not related, and therefore they belong to different subclasses. The orientation of the tRNA substrate would be different. **A**, **B**. The putative bacGlyRS:tRNA complex agrees with the complementary charges on the electrostatic surface (A) and the sequence conservation (B), higher in potential tRNA contact regions. **C**, **D**. The domains involved in tRNA recognition are entirely different in both types of GlyRS. The bacterial tail domain and the eukaryal Ins3 domain, which would contact the tRNA TpsiC loop, are not structural homologs. Neither are the ACBDs homologs. The orientation of both types of GlyRS is the same, relative to the catalytic domain (see the orientation of GSA in bacGlyRS and of ATP-Gly in euk-GlyRS). Remarkably, tRNA’s orientation marks a clear difference between the two types of GlyRS, which belong to different subclasses because they emerged from two different ancestors (Valencia-Sánchez, 2016).

**Figure S7.**
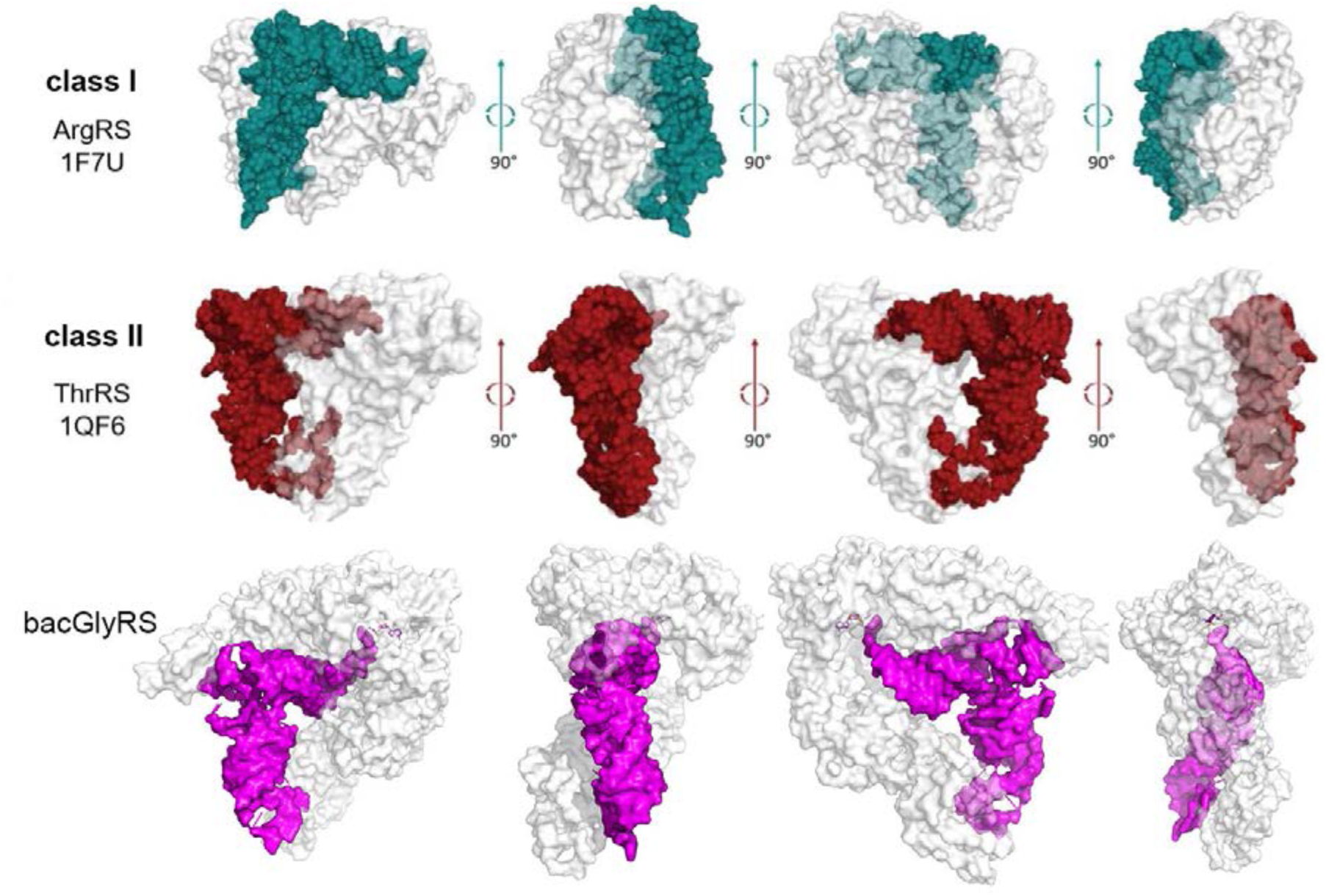
The two types of GlyRS are not related, and therefore they belong to different subclasses. **The interaction of bacGlyRS with tRNA would be a hybrid between class I and II canonical interactions with tRNA.** If our proposed model in complex with tRNA is accurate, the tRNA recognition of bacGlyRS (magenta) would be a hybrid between class I (cyan) and II (red) aaRSs. This model further highlights the differences between the two types of GlyRS.

**Figure S8.**
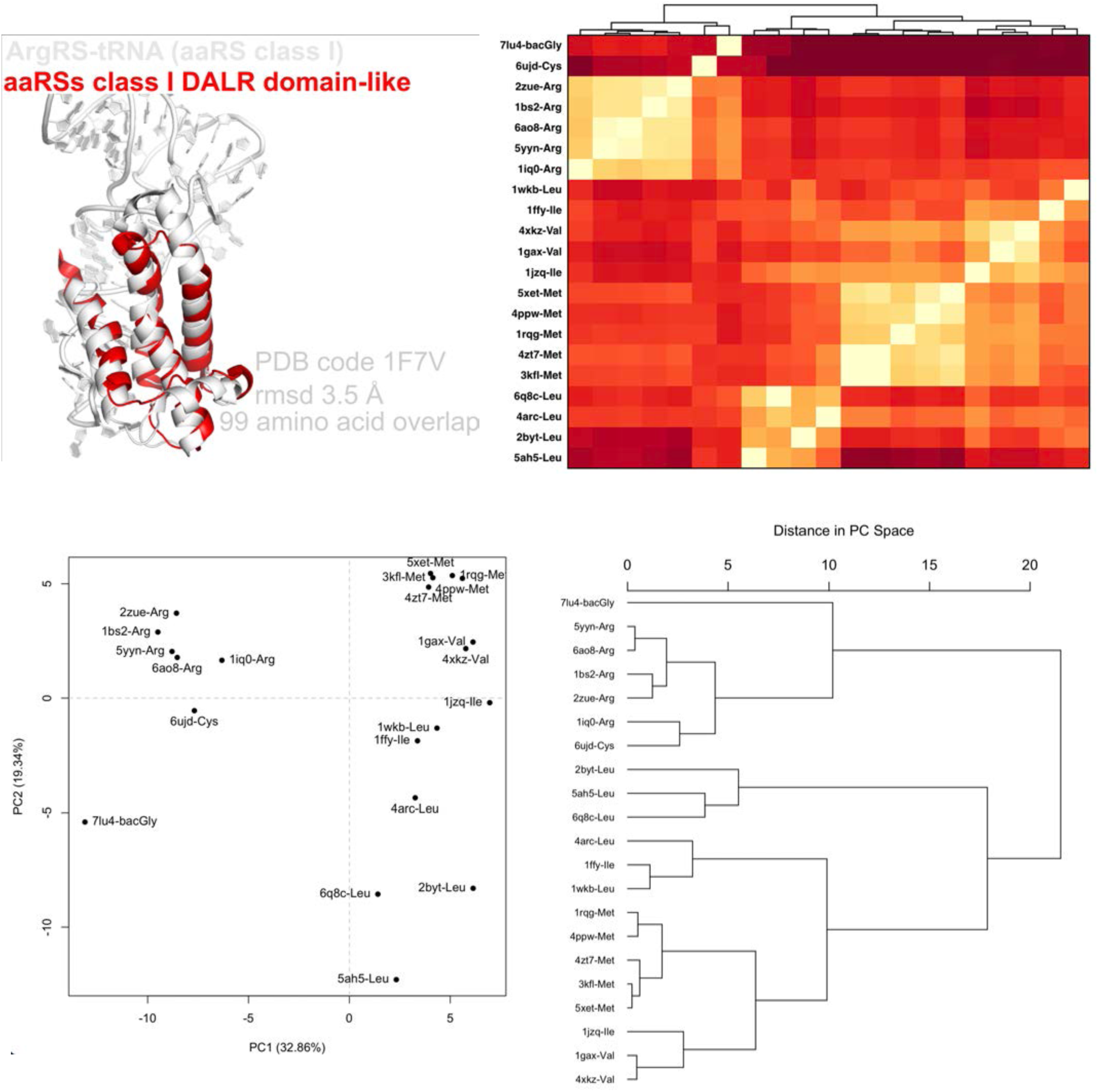
The anticodon binding domain (ACBD) of bacGlyRS is closer to the equivalent region of class I ArgRS and CysRS. The relevant domains of the indicated PDB codes were superposed with the STAMP algorithm on the Multiseq routine of the VMD program (15). The overall RMSD values were analyzed through a Principal Component Analysis (PCA) on Bio3D (16) and BioVinci, and a dendrogram was obtained relating the domains of ArgRS and CysRS to bacGlyRS.

**Figure S9.**
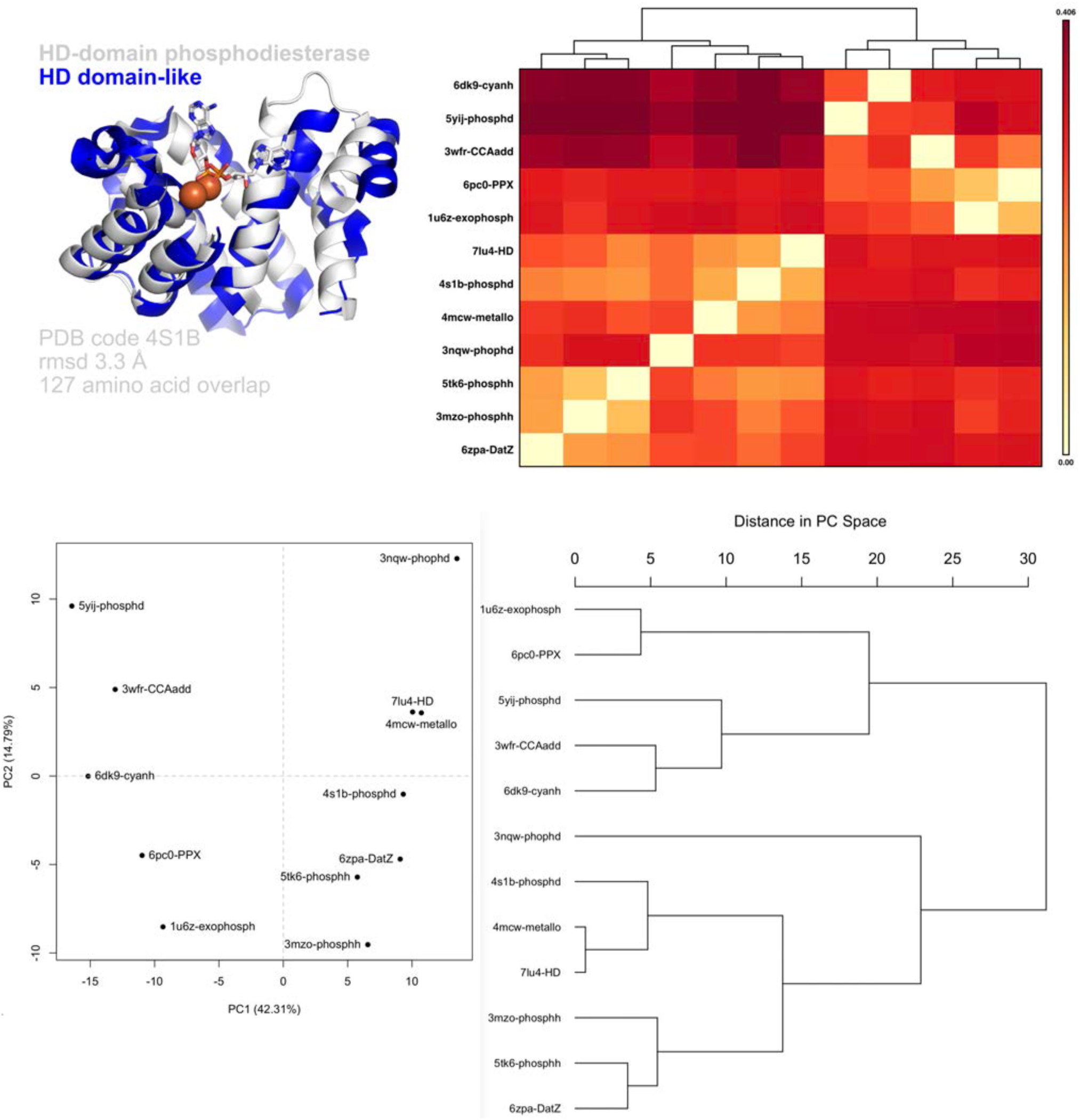
The HD, phosphorylase-like domain of bacGlyRS is closely related to other phosphohydrolases than to proteins with tRNA-related activity domains. The relevant domains of the indicated PDB codes were superposed with the STAMP algorithm on the Multiseq routine of the VMD program (15). The overall RMSD values were analyzed through a Principal Component Analysis (PCA) on Bio3D (16) and BioVinci, and a dendrogram was obtained. At first sight, it was tempting to suggest that bacGlyRS could also have a close relationship with bacterial CCA-adding enzymes of class II (PDB code 3wfr). This analysis shows that apparently, this is not the case. The inactive HD domain of bacGlyRS β-subunit may be instead an ancestor of phosphohydrolases (17). These hydrolases were probably ancestral enzymes in nucleotide metabolism (18)

**Figure S10.**
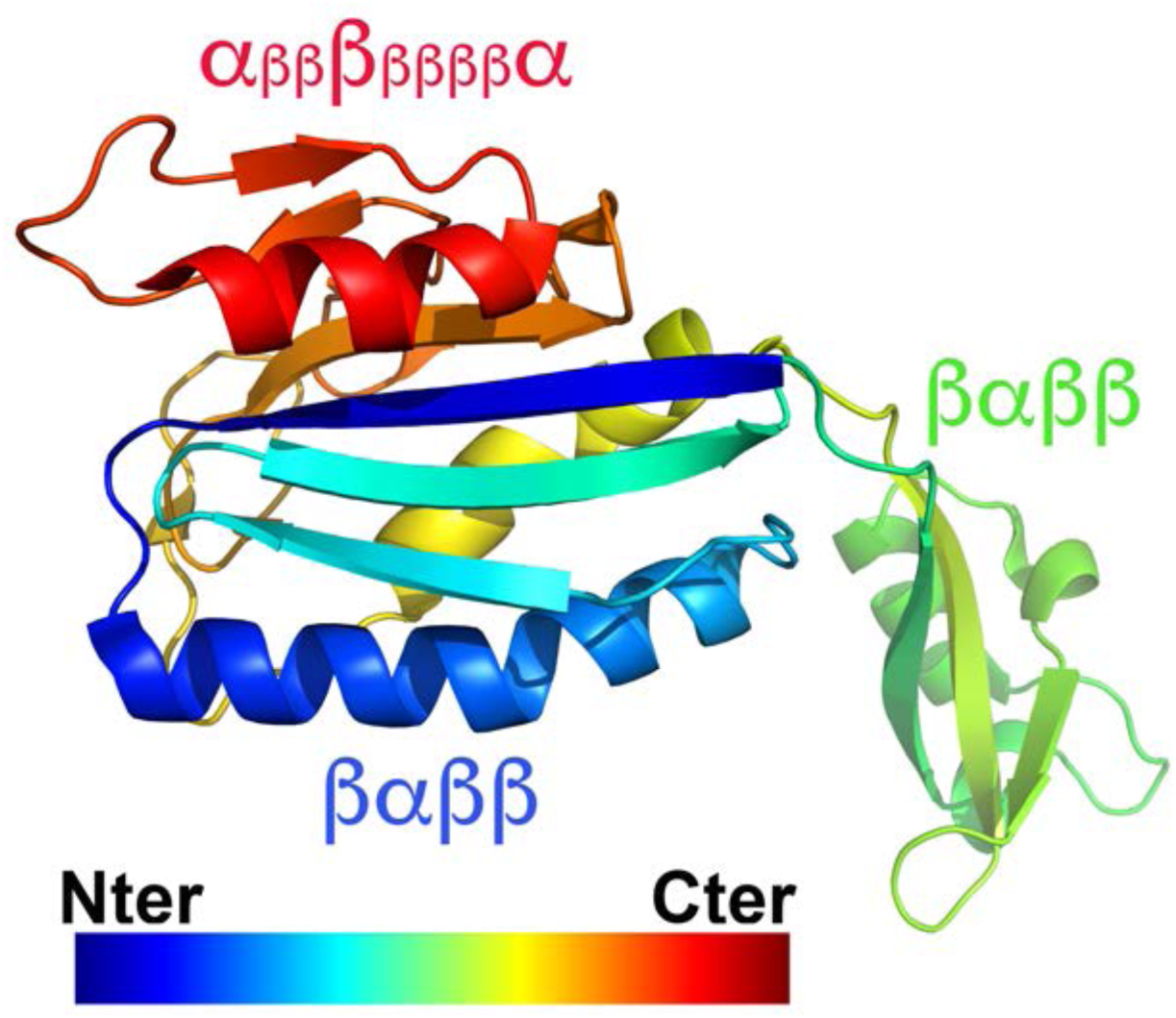
The body domain of bacGlyRS is formed in two parts. In this rainbow-colored illustration from the Nter (blue) to the Cter (red), it can be appreciated that the body domain is wholly formed after the protein chain includes two duplicated βαββ topologies, one of which corresponds to the tail domain. In this sense, the body domain represents a “structural outlier,” as already described in the corresponding domain of the archaeal CCA-adding enzyme (19). Notably, the Cter topology, denoted here as α_ββ_β_ββββ_α(the small caps implicated very short β-strands), has not been described before.

**Figure S11.**
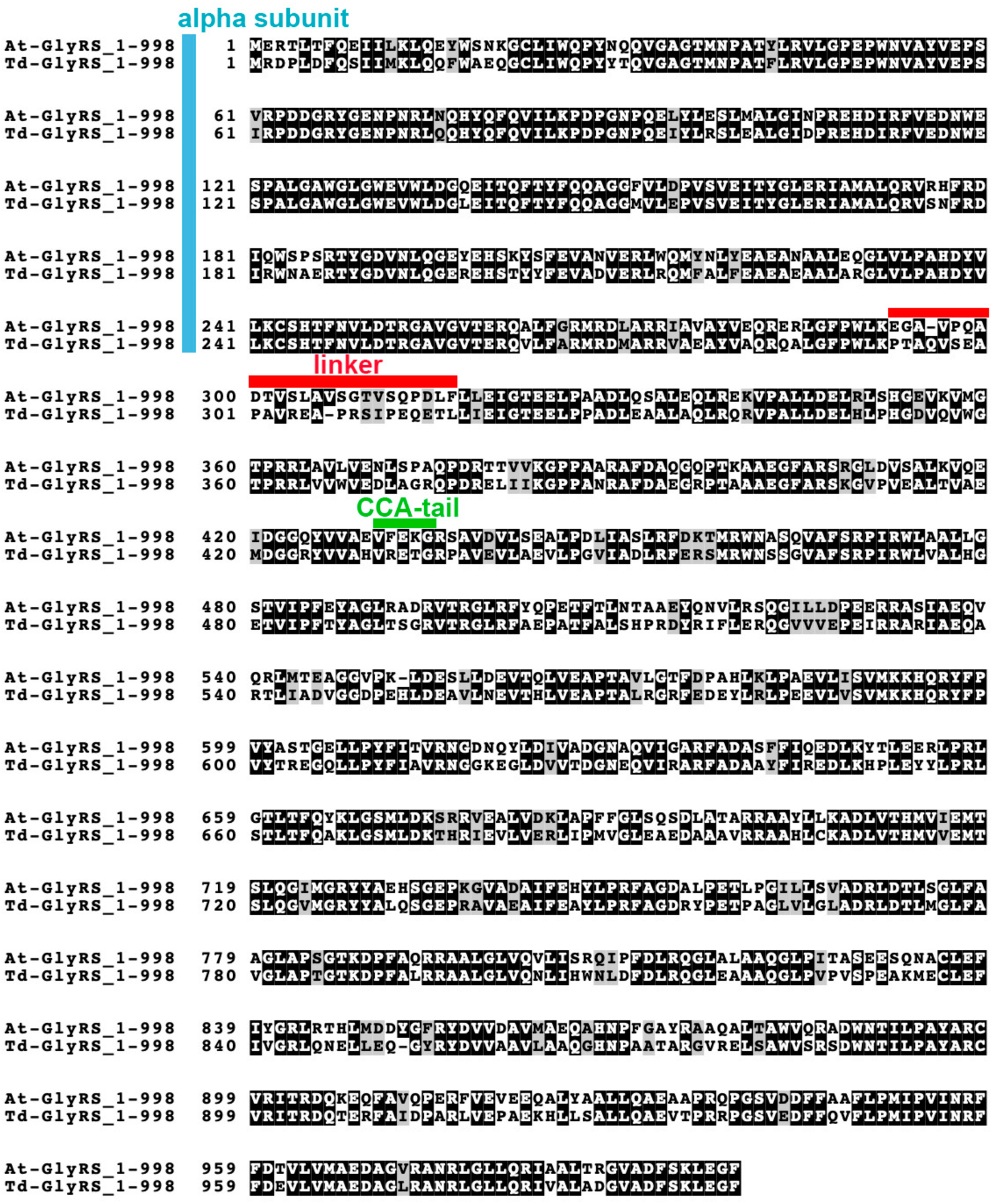
*αβ* At-GlyRS and Td-GlyRS, the two proteins that we use for this work, are highly similar. Sequence alignment of bacterial GlyRS from *Anaerolinea thermophila* (AtGlyRS, 998 amino acids, Uniprot code E8N393) and *Thermanaerotrix daxensis* (TdGlyRS, 998 amino acids, Uniprot code A0A0P6Y0P9). The linker region is found between residues 290-315 (TdGlyRS numbering). The sequences are 71% identical, with 81% similarity. The sites of cleavage, determined by N-terminal Edman sequencing (Alphalyse), correspond to residues TVSQP (309-313 positions, corresponding to the linker region) and VFEKG (430-434, corresponding to the CCA-tail domain) in AtGlyRS. See Figure S12 for the corresponding cleavage sites in the structure. The Figure was prepared using the BoxShade server (https://embnet.vital-it.ch/software/BOX_form.html).

**Figure S12.**
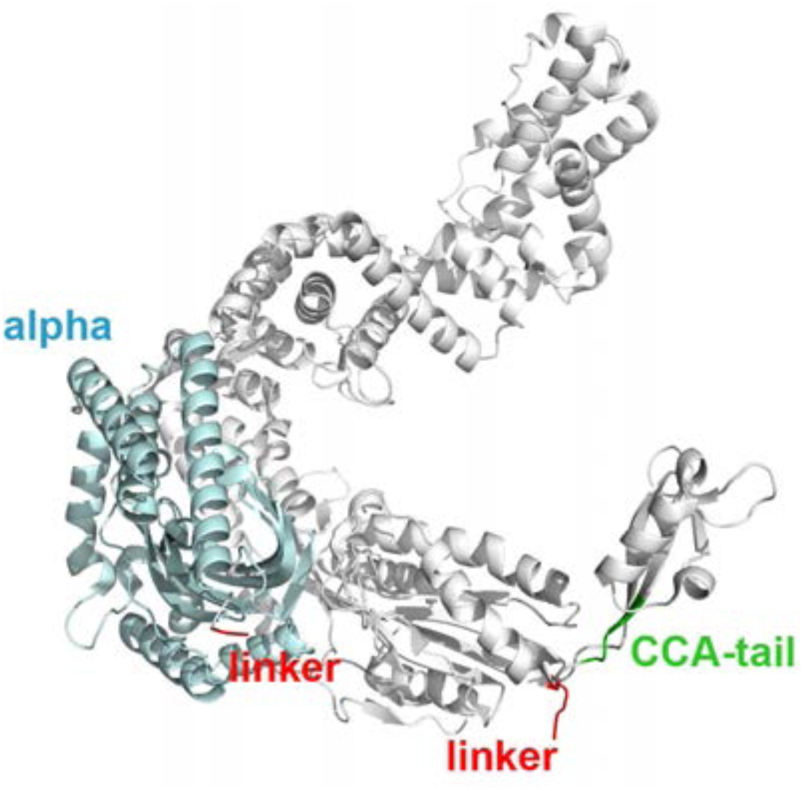
Cleavage sites of AtGlyRS. The structural mapping was done in one *α-b* monomer according to the coordinates of TdGlyRS (this work, PDB code 7LU4). The linker in red is not related to the linker domain (green domain in Figure 2A).

**Figure S13.**
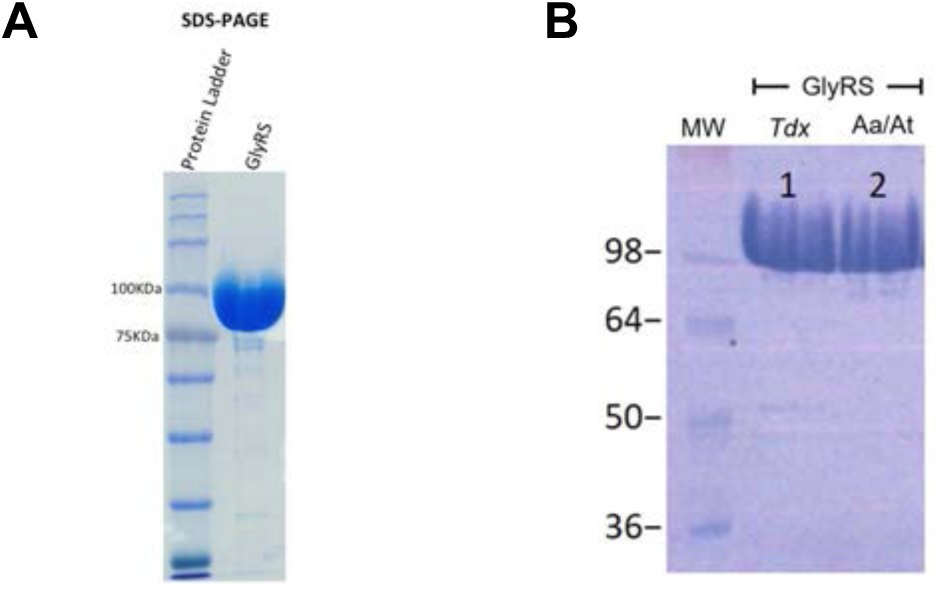
At-GlyRS and Td-GlyRS after purification. **A** 12% polyacrylamide SDS-PAGE showing the purified GlyRS enzyme from *Anaerolinea thermophila* (At-GlyRS, see Supplementary Methods). **B.** Comparison between the purified At-GlyRS and Td-GlyRS.

**Figure S14.**
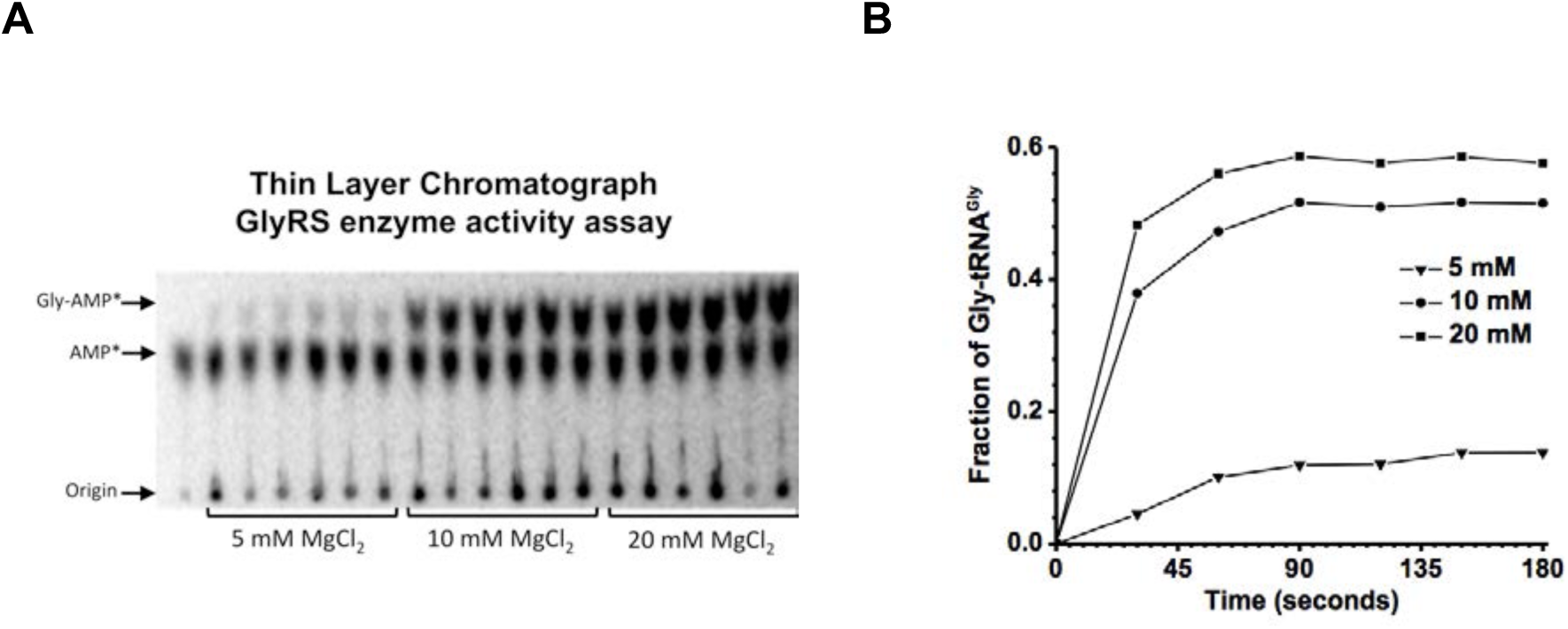
AtGlyRS is active. Aminoacylation reactions with At-GlyRS and At-GlyRS^Gly^_GCC_. **A** Thin Layer Chromatography (TLC) plate showing the time points for reactions containing 5, 10, or 20 mM MgCl_2_. The best aminoacylation fractions were observed while using 20 mM MgCl_2_. **B.** graphic obtained with the densitometry data from the TLC.

**Figure S15.**
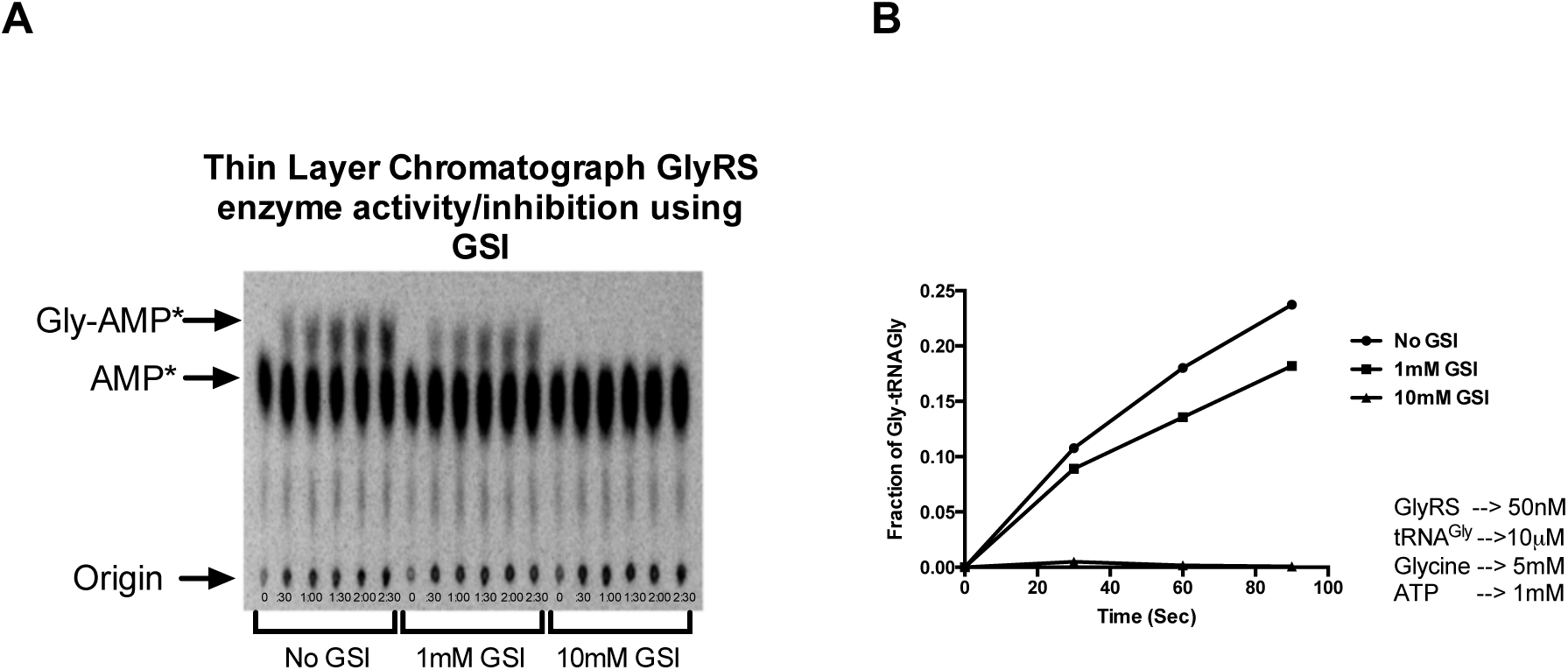
Glycyl-sulfamoyl adenylate (GSA or GSI) inhibits AtGlyRS. Aminoacylation assay was used to verify the effectivity of the GSA analog to inhibit At-GlyRs. **A**. TLC plate showing the time points for reactions without GSI (No GSI), 1 mM GSI and 10 mM GSI. **B**. graphic is obtained with the TLC data; arrows point to the concentrations used for each reagent.

**Figure S16.**
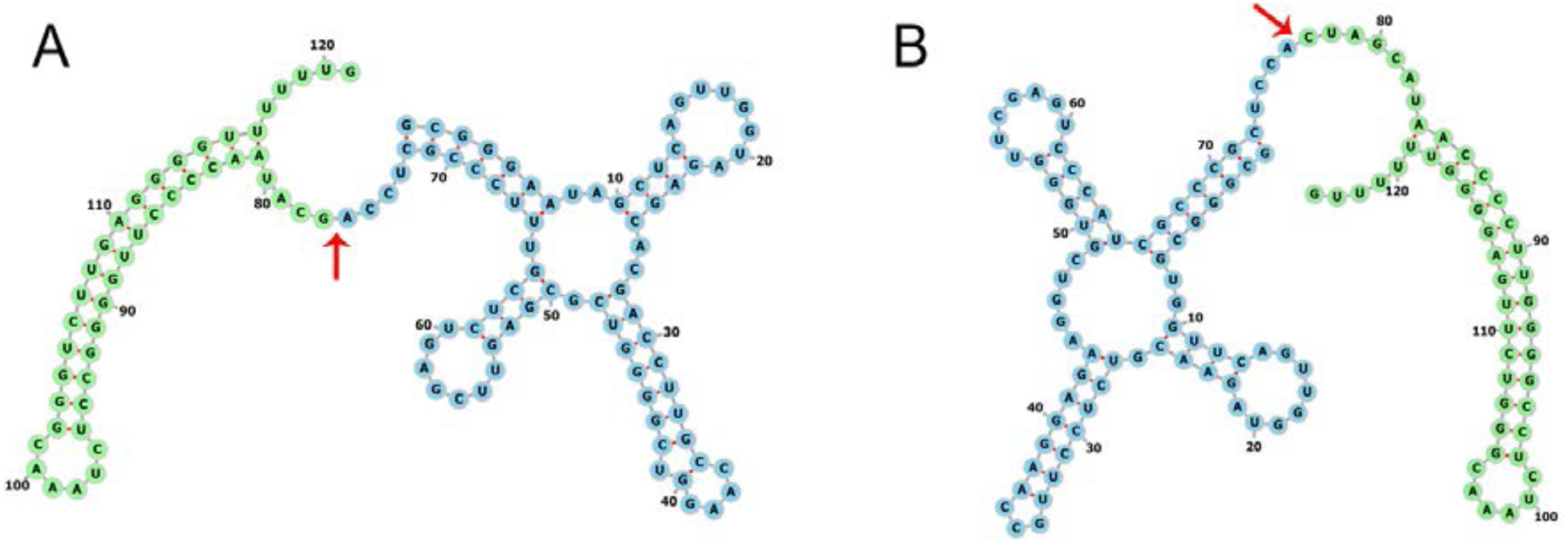
tRNA overexpression on the T7 RNA polymerase system. Representation of the secondary structure of the primary transcripts. Blue, nucleotides belonging to the tRNA sequence; green, nucleotides of the terminator for T7 polymerase. A red arrow indicates the separation between these two elements. **A.** Primary transcript containing the *Escherichia coli* tRNA^Gly^ sequence (GCC anticodon). **B.** Primary transcript containing the tRNA^Gly^ sequence (GCC anticodon) of *Thermanaerothrix daxensis*.

**Figure S17.**
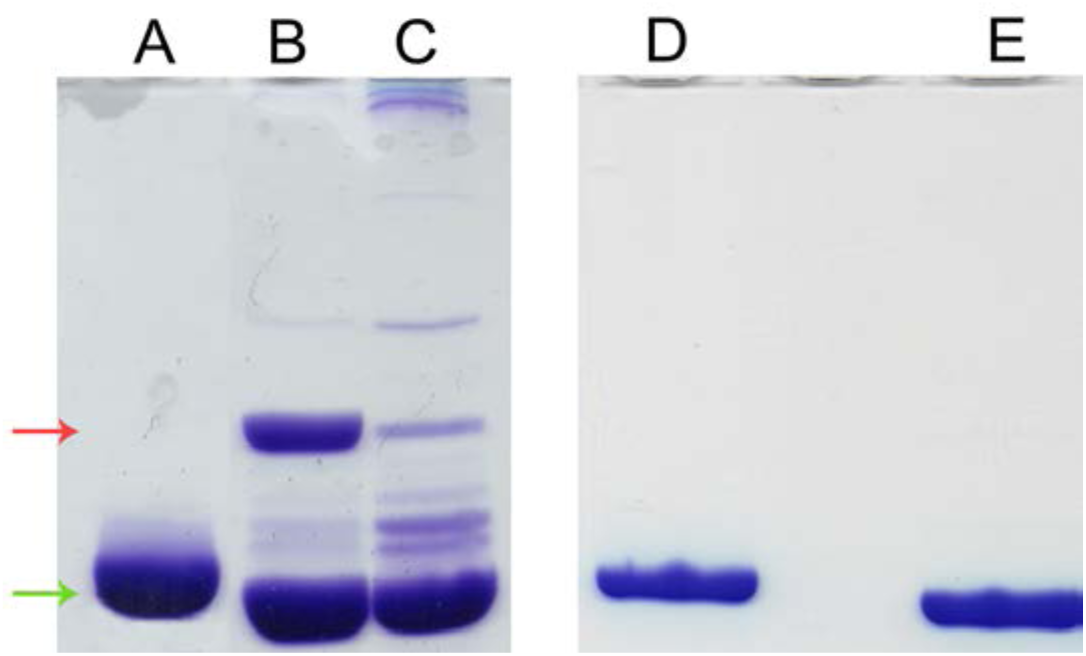
tRNA^Gly can^ be purified by chromatographic techniques. Analysis by UREA-PAGE. On the left side, total soluble RNA of transformed strains and molecular weight control. On the right side, the final purification product is compared with the same molecular weight control. The green arrow highlights the approximate expected weight for a tRNA. The red arrow highlights a second overexpression band. **A** and **D.** Molecular weight control for tRNA. *Aquifex aeolicus*^tRNA Gly^_GCC_ *in vitro* transcription product. **B.** Total soluble RNA of the transformed strain with *E. coli* Gly-GCC tRNA overexpression plasmid. **C.** Total soluble strain RNA transformed with *T. daxensis*tRNA^Gly^_GCC_ overexpression plasmid. **E.** Final purification product with *T. daxensis* Gly GCC tRNA sequence. Bands D and E are expected to have a slight offset due to sequence differences between both tRNAs. This difference gives them slightly different molecular weights.

**Figure S18.**
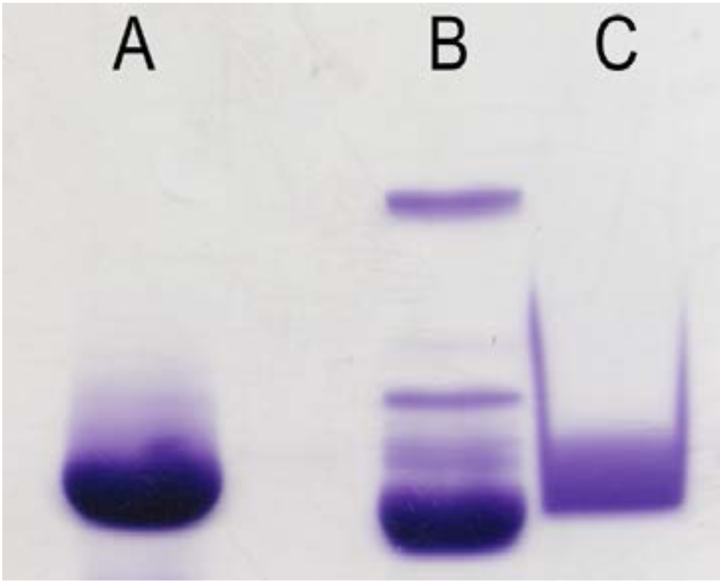
The purified tRNA manages to hybridize with a DNA sequence designed to be complementary to *T. daxensis* tRNA^Gly^_GCC_. UREA-PAGE. **A.** Molecular weight control for tRNA. *Aquifex aeolicus* tRNA^Gly^_GCC_ *in vitro* transcription product. **B.** Total soluble RNA from *E. coli* transformed with *T. daxensis* tRNA^Gly^_GCC_ overexpression plasmid. **C.** Product purification by the hybridization method using a biotinylated DNA oligonucleotide with complementary sequence to *T. daxensis* tRNA^Gly^_GCC_.

**Figure S19.**
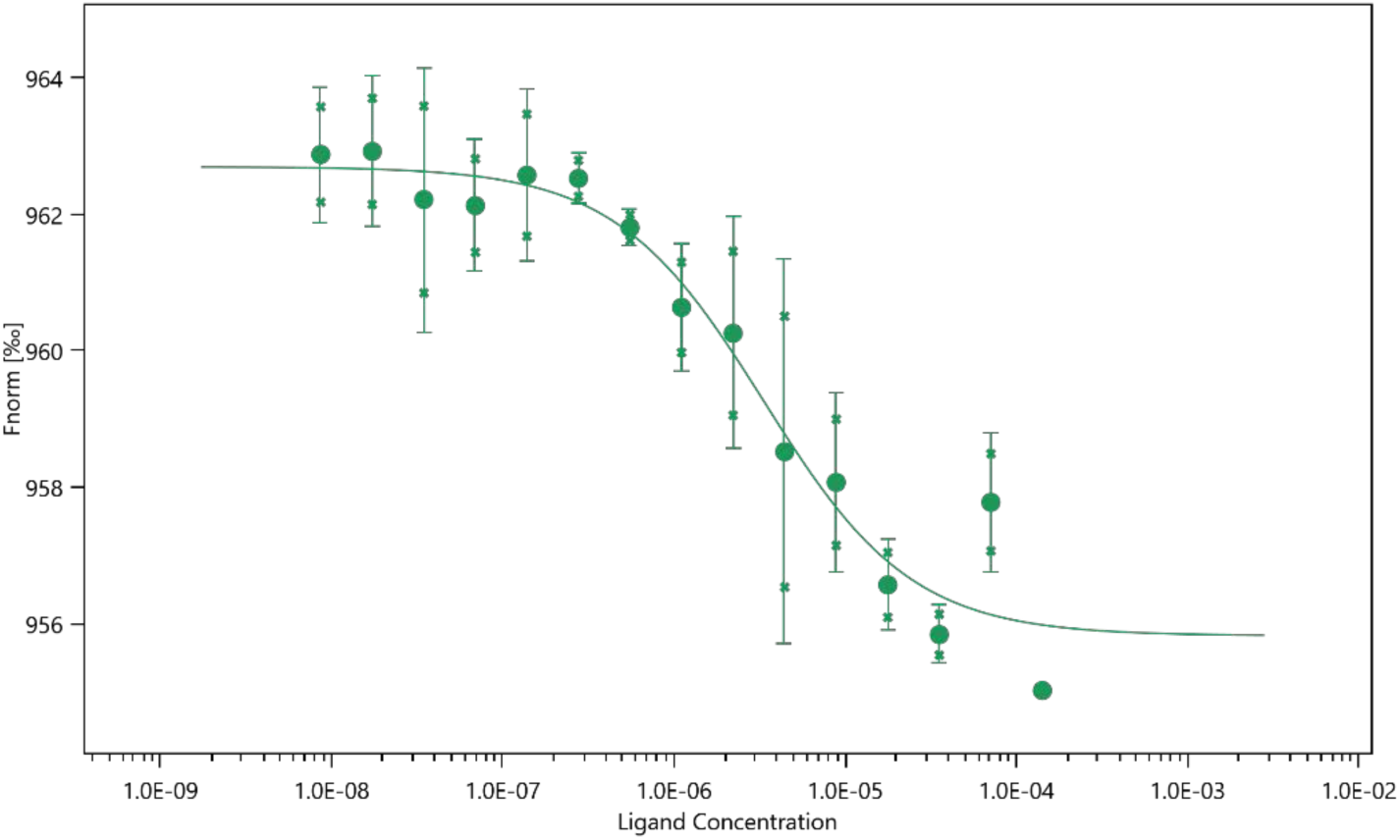
For performing experiments with TDAX protein, a fluorescent label (2^nd^ gen RED) was attached to the protein (His-tag interaction). In this MST experiment, we kept the concentration of labeled TDAX constant at 50 nM, while the concentration of non-labeled tRNA was varied between 142 µM – 8 nM. The assay was performed in 10 mM HEPES pH 7.4. After a short incubation, the samples were loaded into Monolith™ NT.115 Premium Treated Capillaries, and the MST analysis was performed using a Monolith NT.115. Concentrations on the x-axis are plotted in Molar units. A *K_d_* of 3.3 µM was determined for this interaction.

**Figure S20.**
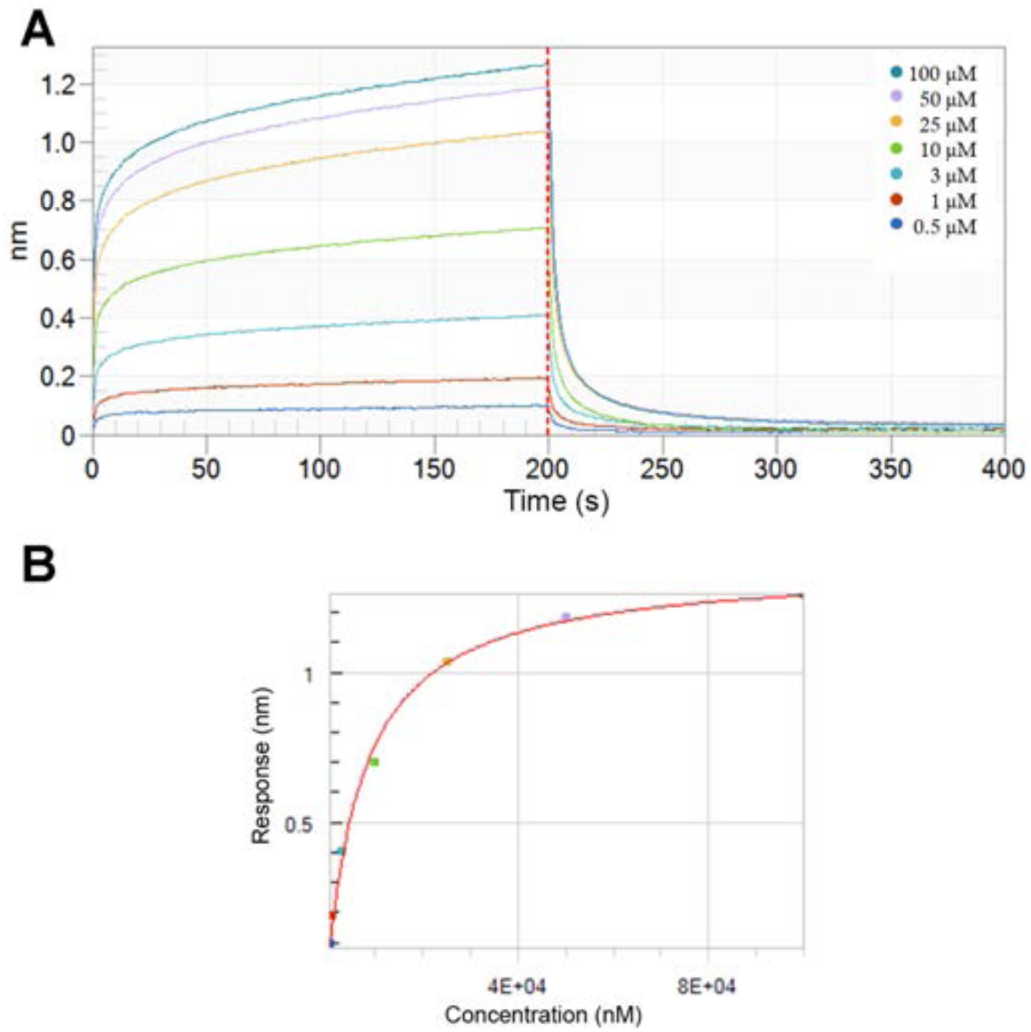
The β subunit accounts for most of the tRNA binding effects in TdGlyRS. Biolayer interferometry binding kinetics assay. **A.** Sensogram shows the association and dissociation sections of the tRNA in the presence of the β-TdGlyRS subunit immobilized to the biosensor. **B.** Steady-state analysis of the data presented in A. The experiment was done in buffer HEPES pH 7.5 50 mM, MgCl_2_ 50mM, ammonium acetate 5 mM, and spermine 5 mM. The K_D_ value obtained using this data is 7.80 ± 0.67 µM (R^2^: 0.995).

**Figure S21.**
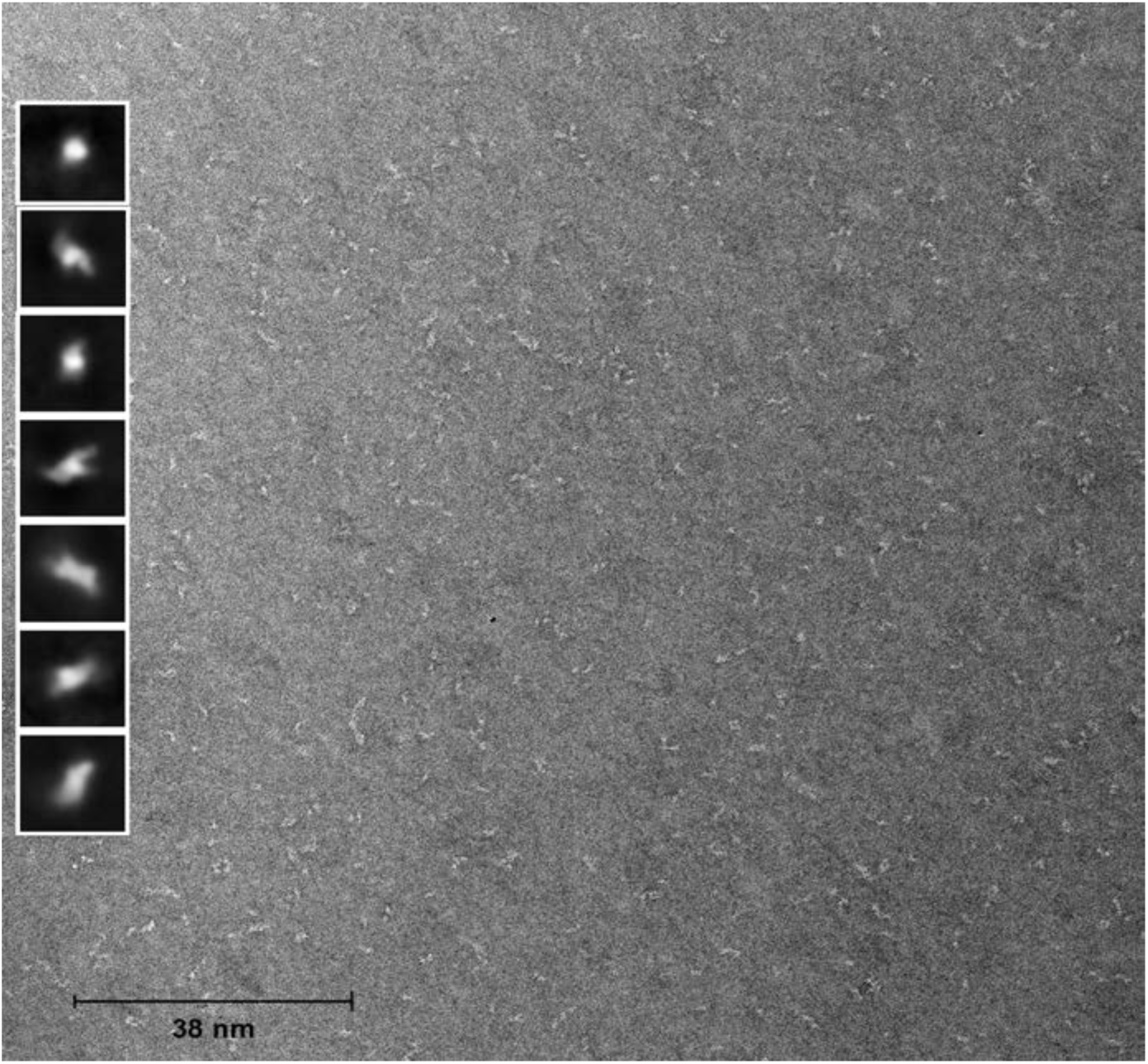
Structure of bacterial GlyRS in solution. Experimental data of negative-stain TEM. Representative example of a micrograph of TdGlyRS^apo^. On the left side are the average two-dimensional classification of the particles according to their orientation. See Table S5 for some parameters of the collected data.

**Figure S22.**
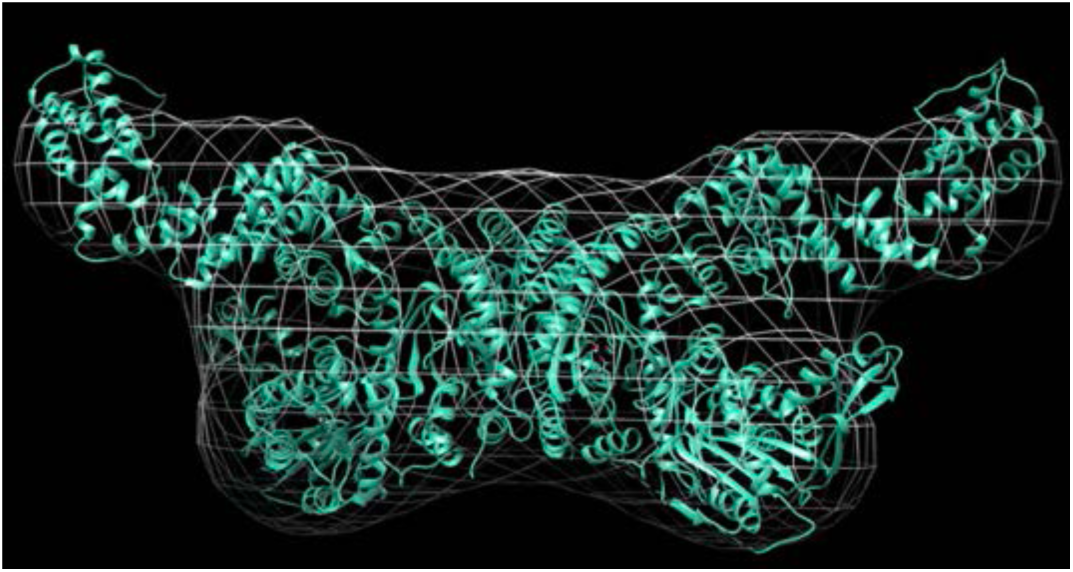
Structure of bacterial GlyRS in solution. 3D reconstruction. Three-dimensional particle reconstruction of bacGlyRS from single-particle electron microscopy data by negative-stain TEM. Overlapping the three-dimensional reconstruction of bacGlyRS by electron microscopy - white mesh - and by X-ray crystallography - cartoon in cyan -.

## Supplementary Tables

**Table S1.** Data collection and refinement statistics

| | $\alpha\beta$-TdGlyRS-Gly |
| --- | --- |
| **Data collection** |  |
| Space group | C2 |
| Cell dimensions |  |
| *a*, *b*, *c* (Å) | 238.58, 127.86, 133.06 |
| $\alpha$, $\beta$, $\gamma$ (°) | 90.0, 103.6, 90.0 |
| Resolution (Å) | 37.62-2.50 (2.54-2.50) * |
| *R*<sub>sym</sub> or *R*<sub>merge</sub> | 0.126 (3.32) |
| *I*/σ*I* | 14.6 (1.5) |
| Completeness (%) | 99.2 (99.1) |
| Redundancy | 17.1 (15.8) |
| Mean (*I*) half-set correlation | 0.999 (0.452) |
| CC<sub>(1/2)</sub> |  |
| Wilson B-factor (Å<sup>2</sup>) | 72.9 |
| **Refinement** |  |
| Resolution (Å) | 35.74-2.50 (2.53-2.50) |
| No. reflections | 132944 (4301) |
| *R*<sub>work</sub>/ *R*<sub>free</sub> | 0.207/0.228<br>(0.363/0.406) |
| No. atoms |  |
| Protein | 14710 |
| Ligand/ion | 5 |
| B-factors |  |
| Protein | 81.0 |
| Ligand/ion | 76.7 |
| R.m.s deviations |  |
| Bond lengths (Å) | 0.004 |
| Bond angles (°) | 0.586 |

One crystal was used.
\*Highest resolution shell is shown in parentheses.

**Table S2.** **bacGlyRS as an ancestral protein**

| Feature | Evidence |
| --- | --- |
| Coevolution with the primordial code (second codon position appeared first-coded in acceptor stem, first codon position seems to be second-coded in anticodon). | $\alpha$: Catalytic domain groups with AlaRS (subclass IId); Ala, as Gly, is a crucial aa defined by the second codon position (Ala, Gly, Asp, Val are coded by the primordial code C, G, A, U).<br>$\beta$: ACBD groups with ArgRS and CysRS, corresponding to amino acids coded by G on the second codon position and C/A, U on the first codon position. |
| tRNA age (acceptor stem first). | Triplicated $\beta\alpha\beta\beta$ (ancient, ferredoxin-like protein topology), with recognition motif for tRNA acceptor, the most ancient tRNA domain. |
| tRNA size (gene duplication event to generate AC stem). | β-subunit seems to have grown as the tRNA became larger (first architecture: $\alpha+\beta$ for acceptor; then $\alpha$ for anticodon). |
| Independent subunits/function (aa-specific and tRNA-promiscuous binding may have evolved with different constraints/timelines). | $\beta\alpha\alpha\beta$ subunit organization (central $\alpha_2$ homodimer, specific aa-tRNA recognition; $\beta$, tRNA recognition by size, not sequence-specific). Analog to (20) |
| Multifunctionality (in LUCA, a few proteins may have performed several related functions). | Body-tail RNA recognition module present in three different functions (aminoacylation, CCA polymerization, and ribosomal processing). |
| Widespread protein domains (the more widespread in organisms and functions, the more ancient). | Present in the three domains of life:<br>α-Core of catalytic domain (all class II aaRSs-catalysis)<br>β-ACBD (seven class I aaRSs-binding)<br>β-Body-tail (three RNA processing enzymes-binding)<br>β-HD-like domain (phosphohydrolases-catalysis) |
| Homology (if the homologs are not easily detected, then the protein may be more ancient). | All domains have borderline sequence similarities (8-13%) with their closest structural neighbors. |
| Non-canonical universal aaRS elements (the less canonical, the less recent). | $\alpha$: An atypical sequence of motifs 1 and 2 on the catalytic domain, leading to non-canonical α-α dimer stabilization and amino group recognition of the cognate amino acid.<br>$\beta$: Atypical orientation of ACBD relative to the catalytic domain, leading to a putative inverted tRNA orientation regarding aaRSs of class II.<br>Hybrid; includes critical elements of both classes of aaRSs;<br>$\beta$: class I AC tRNA recognition; $\alpha$: class II aa recognition. |
| Ancient topology repeats or peptide tandem repeats (proteins are thought to have originated from peptide/motif repeats). | Six α-helices in α-subunit (top of catalytic domain; ensemble for monomer:monomer interactions).<br>Sixteen α-helices in β-subunit (HD-like domain and ACBD).<br>Three $\beta\alpha\beta\beta$ motifs in β-subunit (body-tail-linker domains). |

**Table S3.** **Name and classification of bacGlyRS domains**

| subunit | name | CATH topology | CATH superfamily | SCOPe fold | SCOPe family | Pfam family | ECOD/PASS2.6 |
| --- | --- | --- | --- | --- | --- | --- | --- |
| $\alpha$<br>188-286 | oligom module | Up-down bundle | 1.20.58.180 (PDB 5f5w) | | | PF02091 | |
| $\beta$<br>313-372<br>436-523 | body | Alpha-beta plaits | 3.30.70.590 (PDB 1r89) | Ferredoxin-like | d.58.16.2 | | X: alpha-beta plaits<br>H: (4m5d, 6rxv (U3)), (1q79, 2q66 (polyA)), (1r89 (CCA-add)) |
| $\beta$<br>378-432 | tail | Alpha-beta plaits | 3.30.70.1550 (PDB 1r89) | Ferredoxin-like | d.58.16.2 | | |
| $\beta$<br>526-641 | linker | Alpha-beta plaits | 3.30.70.60 (PDB 2yy3) | | | | |
| $\beta$<br>647-774 | HD-like | Orthogonal bundle | 1.10.3210.10 (PDB 4mcw) | HDdomain/ PDEase-like | a.211.1.1 | PF01966 | Alpha complex topology.<br>X: PDEase-like |
| $\beta$<br>833-992 | ACBD | Orthogonal bundle | 1.10.730.10 (PDB 1bs2) | ACBD aaRSs class I | a.27.1.1 | PF05746 | Alpha bundles<br>X: ACBD class I |

**Table S4.** **αβ GlyRS linker sequences**. Most bacterial GlyRSs are encoded on separate α and β subunits. However, some organisms present a single protein (*αβ*) joined through a linker sequence. In this work, we used two *αβ* fused, highly related sequences from two thermophilic microorganisms. We used the natural linker sequences and a linker sequence from DNA polymerase from bacteriophage RB69 for AtGlyRS. The modification of this sequence has only a moderate influence on the protein’s melting temperature (Tm). See Supplementary Discussion for further details.

| Protein | Linker sequence | $T_m$ |
| --- | --- | --- |
| TdGlyRSwt | WLKPTAQVSEAPAVREAPRSIPEQETL | 73.5 |
| AtGlyRSwt | WLKEGAVPQADTVSLAVSGTVSQPDL | 76 |
| AtGlyRS-RB69 | WLPQGRSHPVQPYPGAFVKEPIP | 69 |

**Table S5.** **Parameters of the collected data of negative-stain TEM on TdGlyRS**. Data collection was performed under a JEOL 1400 microscope operating at 120 kV, equipped with a Gatan 4k x 4k Ultrascan CCD camera. All images were collected at a magnification of X30k automatically by the Leginon program (21). The data was processed in the Relion3 software package by the procedure detailed in a software manual (22).

| <u>TdGlyRS</u> |  |
| --- | --- |
| Number of micrographs | 225 |
| Number of particles | 14,711 |
| Symmetry <sup>a</sup> | C2 |
| Resolution Limit (Å) | 21.85 |
| Pixel Size (Å) | 3.8 |
| Box size (pixels) <sup>b</sup> | 92 |
| Number of classes -2D- | 7 |
| Mask size (Å) <sup>c</sup> | 340 |

<sup>a</sup> This parameter refers to the symmetry imposed for the generation of three-dimensional reconstruction.
<sup>b</sup> Size designated to the box for particle extraction step.
<sup>c</sup> For the generation of the three-dimensional model.

## Supplementary Results

We obtained highly purified, naturally linked *αβ*-subunits of GlyRS of *Thermoanaerotrix daxensis* (TdGlyRS, Figures S9-S11) using a His-tag affinity chromatography step and a heat treatment. Negative-staining electron microscopy confirmed the ensemble’s dimeric nature and relative orientation (Figures S21-22, Table S5). TdGlyRS binds tRNA^Gly^, with an affinity of 3 μM (Figure S19). Using a highly similar protein to TdGlyRS from *Anaerolinea thermophila* (AtGlyRS), we verified that the whole, linked enzyme could perform the aminoacylation reaction (Figure S14). The reaction was inhibited by a transition state analog, glycyl-sulfamoyl adenylate (GSA) (Figure S15). These findings provide biological relevance to our reported crystal structure.

To discover the architecture of the whole enzyme and to unveil the folding domains of the β-subunit, we solved by molecular replacement using the coordinates of the α-subunit dimer and automatic building, the crystal structure of TdGlyRS in complex with glycine in one monomer at 2.7 Å resolution (Table S1). The electron density map (Figure S2) unambiguously showed all features and domains of the 200 kDa enzyme and the bound glycine in one monomer of the biological dimer found in the asymmetric unit (Figure S3B).

As previously shown (PDB entry 5F5W), the α-subunit forms a homodimer that also functions as the dimeric interface of the α_2_β_2_ ensemble (Figure 1), with 58% of its interface created by a 97 residue α-helical region located at the C-terminal part of the subunit and situated on top of the signature antiparallel β-strand of class II synthetases (Figure S3B). There is no conformational change in the α_2_ homodimer in complex with the β-subunit (PDB entry 7LU4) compared to the structure of the α_2_ homodimer alone (PDB entry 5F5W), (Figure S3B). Surprisingly, the β-subunit does not participate in the stabilization of the heterodimer interface. Instead, each β-subunit only interacts with one α-subunit. The α-β interface is formed mainly by highly conserved residues (Figure S4B) complementary in shape and charge (Figure S4B, C), which further supports the geometry of their interaction.

The comparison of the apo monomer of TdGlyRS with the glycine-bound monomer (Figure S3B) reveals the same conformational changes already reported for the crystal structure of the α-subunit of *Aquifex aeolicus* in complex with an analog of glycine-adenylate (GSA or G5A); a ∼5 Å displacement of an 11-residue region where a Trp forms a cation-π interaction with glycine (Figure S3B). Thus, the reported crystal structure bacGlyRS represents a productive, biologically meaningful complex. The presence of glycine bound and the correlated conformational movement already reported for the binding of GlyAMS (14) shows that the binding of the amino acid alone is enough to promote this movement and that this conserved region, shared with AlaRS, is important for the recognition of the amino acid.

## Supplementary Discussion

### - The protein size of bacGlyRS is counterintuitive

In general, the lengths of orthologous protein families in Eukarya almost duplicate the sizes found in Bacteria and Archaea (see referenced work in (23)). Regarding aaRSs, “most of their additional domains, seen as extensions or insertions, with the exception of the ACBD and editing domains, are present in higher eukaryotes and completely absent in the bacterial domain, which makes any aaRS larger as the tree of life ascends” (24).

The size of bacGlyRS is counterintuitive because it is much bigger and has more domains than eukGlyRS (Figure S5). bacGlyRS represents a very long protein anomalous in bacteria. Remarkably, bacGlyRS is present even in bacteria with a predicted meager number of multidomain proteins, like *Chlamydia psittaci* (25).

Regarding aaRSs domains, “the extra domains possessed by aaRSs are mostly particular to these enzymes and not found elsewhere, e.g., the CP1 editing domain; the Add1 domain of ArgRS, the INS domain of ProRS, leucine zippers, glutathione S-transferase (GST) domains, WHEP and EMAPII domains. Many of the aaRSs extra domains specific to a particular aaRS do not have homology to other proteins” (26).

Taking together this information with the data presented in this work, bacGlyRS shows signs of a possible “frozen” evolutionary accident that allows analyzing a molecular fossil in the present day.

### - The α and β subunits of bacGlyRS dissociate and cleave easily

We tried to crystallize several bacterial GlyRSs expressing separated α and β subunits from different sequence sources. Most of them dissociate the subunits in the crystallization drops. An alternative strategy was to look for an enzyme with naturally fused α and β subunits from a thermophilic organism, *Anaerolinea thermophila* (Figures S11-15). However, it was observed that the α and β subunits were cleaved on the linker region during the purification protocol (Figures S11-S12). Several natural linkers were substituted on the sequence to prevent cleavage. A linker derived from the DNA polymerase sequence of bacteriophage RB69 (Table S4) joined both subunits to obtain a chimeric protein of *A. thermophila,* which is active (Figures S14 and S15) and formed crystals that diffracted at poor resolution.

We explored the crystallization properties of proteins very similar to that of *A. thermophila.* In particular, from the sequence *of Thermanaerothrix daxensis,* which bears 70% sequence similarity to that of *A. thermophila* (Figure S11). We obtained crystals that diffracted at very low resolution, but we improved them using different optimization techniques (see Supplementary Methods).

### - Bacterial GlyRS is at the origin of all aaRSs and non-sequence specific RNA binding activities

Among aaRSs, bacGlyRS represents a unique α_2_β_2_ oligomeric arrangement. One of the problems that we faced during the development of this project is that the α and β subunits dissociate easily. The α-subunit of bacGlyRS is the only known case among aaRSs where the catalytic domain is not covalently attached to other domains. The structure shows the extent of this relative independence, as the α-subunit is exclusively responsible for the tetrameric assembly. It forms the same α_2_ homodimer that can perform the first step of aminoacylation (14). The α-subunit seems an entity that may have evolved unconstrained from the β-subunit. In contrast, the β-subunit may reflect the restraint coevolution with a tRNA substrate that became bigger with time. The presence of two different protein architectures (α and α+*b*) in distinct potential tRNA recognition stems on the β subunit might also indicate a two-stage evolution scenario.

The ArgRS-like, bacGlyRS ACBD represents a connection between class I and II aaRSs. Besides bacGlyRS, no other class II aaRS has another critical domain shared with class I aaRSs. Regarding a linkage with the genetic code (Figure 5), remarkably, bacGlyRS, ArgRS, and CysRS are clustered in the same group according to their ACBD (Figure S8, Figure 5).

Thus, although our evolutionary proposal (see below) implicates that the ACBD was a later addition to bacGlyRS, the ACBD incorporation should have happened very early during the development of the genetic code. The ACBD widely spread in most class I aaRSs, probably represents the most ancient domain of this type (Figure 5).

Regarding the tail-body domains, the sequence similarity of the CCA-adding enzyme of class I with the equivalent domains of the bacGlyRS is so low that, at first glance, it is tempting to speculate that they are not homologs but analogs. However, there are clear signs of homology in this region: two structural units (domains), not only one, are involved in the comparison; the arrangement and the relative orientation of both domains are the same in both proteins; the connectivity of the secondary structure elements is the same; finally, both proteins share the insertion of large parts of secondary structures to form the body domain (PDB code 1R89). Remarkably, the homology with the tail-body domains is also present with proteins involved in eukaryotic ribosome processing (PDB code 4M5D). Thus, this region is widespread in the three life domains and connects with general RNA metabolism.

As for the linker domain, although there are issues to find a clear homolog of it, we identified a βαββ topology that is also found in the body domain in tandem with the tail domain; this raises the possibility that a triplication event is the origin of the *αβ* architecture of bacGlyRS. Thus, a repeat could be created as it was required. As the sequence conservation between the tail-body-linker domains is extremely low, the structural motifs highly degenerate, and the domains are embellished with other structural motifs, this triplication, if true, must have taken place relatively early in evolution. Notably, the βαββ topology found in these domains represents a ferredoxin-like fold, which is considered ancient and central to the origin of metabolism.

Concerning the evolution of bacGlyRS, the alternative hypothesis to the one stated in the Discussion would be a relatively recent appearance of the bacGlyRS, requiring the development of an entirely novel, multidomain protein. Another possibility would be the HGT of the *αβ* architecture from archaea, with an atypical tRNA recognition, provided that the canonical tRNA interaction pattern was already S-29 present in class II aaRSs, and then the concomitant loss of the much shorter, single-chain eukGlyRS. This hypothesis seems less likely in the current structural scenario.

Thus, regarding bacGlyRS, our work defines the structure of the entire bacterial GlyRS α_2_β_2_ heterotetramer, thoroughly delineates the differences between the two types of GlyRS, proposes a model of the tRNA recognition process, and provides insight that poses α_2_β_2_ GlyRS at a landmark on the origin of aaRSs and other proteins involved in RNA metabolism, thus offering perhaps the first clear example of the coevolution of the Ribonucleoproteic (RNP) world. Further work is needed to corroborate the tRNA interactions and the chemical details of the aminoacylation reaction.

### - The rooted bacterial Tree of Life (ToL) can be inferred from ancestral topologies

In general, performing a phylogenetic analysis with any aaRS is very complicated because of the homologies with other aaRSs. Correctly resolving orthologs, paralogs, and horizontal gene transfer (HGT) is very difficult to solve. bacGlyRS offers a unique window to analyze these ancient proteins because due to its localized nature in bacterial species, the sequences selected correspond to unambiguous orthologous genes. There are several choices to perform a phylogenetic analysis on bacGlyRS: 1) on the α-subunit, catalytic domain; the interactions of aaRSs with the cognate amino acid and ATP occur in this domain. Two different folds of this domain define the two classes of aaRSs, I and II (27). This class-defining domain is considered the ancient tRNA synthetase, and bacGlyRS has the shortest sequence among the class II aaRSs (28). Thus, the bacGlyRS catalytic domain is the simplest by far among aaRSs. Moreover, the helical repeats that cover it may represent primordial oligomerization topologies. In addition, the evolution rate of the whole α-subunit may be completely independent of the β-subunit. Indeed, the α-subunit is much more conserved, indicating that the β-subunit is more ancient.

The β-subunit presents two types of multidomain organizations based on two of the most basal folds: α and α+*b*, both of which may have originated from multiple duplication events (29). Or we may choose to analyze 2) the α-helical, anticodon binding domain (ACBD). Regarding the interactions of aaRSs with tRNA, these are mainly done in specific ACBDs. There are three different ACBDs known, being the most expanded the α-helical domain of class I aaRSs. Interestingly, ArgRS, which ACBD is the closest related to bacGlyRS, is the most deeply rooted class I enzyme (26). Option 3) the duplicated βαββ ancient protein topology, which in principle sits the ancient tRNA TpsiC-loop (this work and (30)), which sequence is highly conserved and defines the limit of the acceptor stem (reviewed in (31)).

Most known, currently accepted phylogenies are made with either a single or few genes from whole, multidomain genes/proteins (32) or using phylogenomics approaches (10), (9). These approaches do not consider the different evolving rates of protein domains and protein domain segments. HGT and orthology determination biases are also common in bacterial phylogenies (33). With our proposed approach, we believe that several of these issues may be overcome.

Surprisingly, the derived tree agrees very well with several available trees and other available evidence. It also supports previously suggested phyla relationships and even places the root very close to Fusobacteria, recently proposed as the most ancestral phyla (9). Thus, we believe that this classification is supported by our data and several previously scattered pieces of evidence. We recommend dividing the bacteria domain into four superphyla according to the obtained tree (Figure 4): Desulfophyla, Firmicutes, Cyanobacteryota, and Proteobacteria.

## Supplementary Methods

### Structural and sequence analysis

We used a combination of the outputs of both the Dali Server (http://ekhidna2.biocenter.helsinki.fi/dali/ (34) and DASH (Database of Aligned Structural Homologs https://sysimm.org/dash/) (35)) to look for structural homologs of the five domains of the bacGlyRS β-subunit. We prepared a collection of the trimmed models (PDB codes) comprising only the relevant possible homolog domains, and we calculated an overall multiple structural alignment using the STAMP algorithm (36) as implemented in MultiSeq in VMD (37). The aligned structures were used in Bio3D (16) to define an invariant core with minimal structural variance among all the protein structures. This core of Cα atoms was the basis for the RMSD matrix (plotted in BioVinci), dendrogram calculation, and Principal Component Analysis (PCA). The structures were superimposed onto the core, and the variances shown by PC1 and PC2 were used to define different subgroups.

For the cases of the body-tail domains of bacGlyRS and archaeal CCA-adding enzyme, we performed structure-based multiple sequence alignments. All the sequences obtained from a BLAST Protein search on the refseq_select database of the body-tail domains were analyzed. The output sequences were aligned with the ClustalW plug-in in MultiSeq. A profile alignment was calculated considering the structural alignment of corresponding domains of the models (PDB codes 1R89, residues 277-379 and 7LU4, residues 316-432). We used IQ-TREE 2 (38), which is suggested to find the highest likelihood scores among ML trees (39). The sequences, log-file with the parameters used, the derived tree, and the file used to obtain the color code figure in iToL (11) are available in the Supplementary Material of the BioRXIV preprint server: https://www.biorxiv.org/content/10.1101/2021.08.20.456953v5.supplementary-material.

### Expression and purification of *Thermanaerothrix daxensis* GlyRS (TdGlyRS)

The plasmid containing the coding sequence of *αβ*TdGlyRS, with the Cys residues mutated to Ser to avoid protein aggregation and a TEV protease cleavage site-His_6_ tag at the carboxyl terminus, was ordered from the company Atum DNA2.0, with the sequence optimized for production in *Escherichia coli*. The protein was purified by affinity to Ni^2+^ and heat treatment at 72°C for 30 min. Crystals of *αβ*TdGlyRS were obtained by sitting-drop vapor diffusion. We obtained a highly reproducible crystallization condition, with very nice crystals that diffracted poorly (control condition; 320 mM ammonium phosphate, 1% dextran sodium sulfate Mr 4,000 (Molecular Dimensions), and 100 mM glycine pH 8.5). These crystals grew S-33 within 4 hours, and we used them in a cross-seeding protocol to obtain crystals from a condition difficult to reproduce: we collected the crystals from 5 to 7 wells in 50 uL of mother liquor and crushed them by vortexing for 2 minutes in the presence of a glass bead. We verified the crushing on the optical microscope, and the seeds were diluted from 10 to up to 10 million-fold.

The crystals (see Methods) were dehydrated for one day, substituting the mother liquor with the cryoprotective solution, which was prepared by mixing 625 μL sucrose at 70%, 200 μL magnesium formate 1M, 50 μL sodium acetate 1M pH 4.6, 25 μL Gly 2 M pH 8.5 and 100 μL dextran sodium sulfate Mr 5,000 at 30%.

### Expression and purification of *Anaerolinea thermophila* GlyRS (AtGlyRS)

BL21(DE3) *E. coli* cells were transformed with GlyRS from *Anaerolinea thermophila* (AtGlyRS) without cysteine residues, with a 6x His tag on the C-terminal and containing a linker region between the alpha and beta subunits corresponding to the one seen in the structure of the bacteriophage RB69 DNA polymerase (PDB code 1WAF; Table S4) (40). The expression plasmid (At-pJ401), with a modified nucleotide sequence for optimal expression in *Escherichia coli*, was purchased from DNA2.0 (Newark, California, USA). After transformation, a colony was selected from agar plates and inoculated a 50 mL LB overnight stock. This culture grew at 37°C on a shaker at 200 rpm for 16 h and was used the next day to inoculate fresh LB-media in a 1:1000 ratio. Five 1000 mL cultures were allowed to grow inside 2 L flasks at 37°C until OD_600nm_ reached 0.6. Protein expression was induced by adding 1 mM IPTG, and cells were allowed to grow 4 hours more under the same conditions. Cells were recovered by centrifugation at 6000 rpm, pellets collected, and frozen at -20°C until needed. For purification, a cell pellet of 10 g was suspended in 100 mL of buffer A (100 mM potassium phosphate pH 7.3, 300 mM NaCl, 5% glycerol). Cells were disrupted by sonication on ice, and lysates were clarified by centrifugation at 12 000 rpm. Cleared lysate was loaded onto a 5 mL Ni-NTA column (GE Healthcare) pre-equilibrated with buffer A and mounted in an FPLC ÄKTA system. The column was washed with 120 mL of buffer A, followed by 80 mL wash with 4 % buffer B (identical to buffer A but with 500 mM imidazole). Protein was eluted with a 20 mL gradient from 4%-100% buffer B. Fractions of 1.0 mL were collected. The ones with higher absorbance at 280 nm (A_280_) were pulled together and precipitated using enough ammonium sulfate salt to achieve a final concentration of 2.0 M. The salt was entirely dissolved by gentle agitation. The suspension was loaded in a 25 mL bead toyopearl-butyl 650 column pre-equilibrated with 100 mM Tris-HCl pH 8.8, 2 M ammonium sulfate and 50 mM NaCl, buffer C. After loading, the column was washed with 30% C; GlyRS was eluted from the butyl column by a gradient from 30 to 0% C. Finally, the protein was dialyzed and concentrated to 6 mg/mL in 100 mM Tris-HCl pH 8.0 and 50 mM NaCl, and 0.5 mL of the protein solution loaded to a pre-equilibrated superdex 200 10/300 GL size exclusion column (GE Healthcare). A single peak was recovered. The SEC procedure was repeated until all the protein was passed. Purity was >95%, judging from SDS-PAGE (Figure S13A). All buffers used contained 0.1 mM PMSF. Lysis buffer was also supplemented with EDTA-free protease inhibitor tablets (Roche).

### Preparation of *Anaerolinea thermophila* tRNA^Gly^ (At-tRNA^Gly^_GCC_) for aminoacylation assays

*Anaerolinea thermophila* tRNA^Gly^_GCC_ was in vitro transcribed as described previously (41). DNA oligos for Klenow reaction were 5’-AATTCCTGCAGTAATACGACTCACTATAGCGGGCGTGGTTCAGTTGGTAGAATGCTT -3’ and 5’-TGGAGCGGGCGATGGGACTCGAACCCACGACCTCTTCCTTGGCAAGGAAGCATTCTACC -3’ and contained no 2’OH modifications as the oligos in the original protocol. The DNA template was completed using the large Klenow fragment (New England Biolabs Cat. No. M0210). The reaction mix of 500 μL contained 4 μM of each oligo, 2.5 mM of each dNTP, 1x NEB buffer, and 5 μL of the Klenow enzyme. The reaction was carefully mixed and divided into 5 PCR tubes. The polymerization reaction was carried out in a thermocycler with 15 cycles of annealing at 10°C for 45 s and a slow ramp to 37°C for 45 sec to allow DNA polymerization. The extended double-stranded DNA template was recovered by 30 min centrifugation at 14,000 rpm after 16 h of precipitation with 2.5× volumes of ethanol, 0.3 M sodium acetate, and 0.2 M acetic acid. The DNA pellet was air-dried and suspended in 200 μL DEPC-treated water. DNA extension was verified using 8M PAGE mini gels.

For a 2 mL transcription reaction, all 200 μL of DNA template prepared from the Klenow extension were used. The reaction proceeded in the presence of 40 mM Tris-HCl pH 8.0, 4 μM spermidine, 25 mM MgCl_2_, 40 mM DTT, 0.01% Triton 100, 4 mM of each NTP, 0.2 mg/mL T7 RNA polymerase, at 37°C for 14 hours. After transcription, the tRNA^Gly^_GCC_ was purified by PAGE extraction, recovered in 10 mM Tris-HCl pH 8.0, 1 mM EDTA (TE buffer), and stored at -20°C until needed.

### At-tRNA^Gly^_GCC_ folding for aminoacylation assays

For aminoacylation reactions, an aliquot of the in vitro transcribed At-tRNA^Gly^_GCC_ was folded before use. To fold the tRNA^Gly^_GCC_, an aliquot at a concentration between 200-400 μM was heated to 80°C for 3 min on a heat block; immediately after, 1/10 of the final volume of an MgCl_2_ solution was added to give 10 mM in the final mix. Then, the heat block was taken off the heat and incubated on the bench to reach ambient temperature. This process takes about 1 h 20 min. After which, the tRNA^Gly^_GCC_ was folded.

### At-tRNA^Gly^_GCC_ labeling for aminoacylation assays

To verify the activity of AtGlyRS, we followed the aminoacylation reaction labeling the in vitro transcribed tRNA^Gly^_GCC_ using a previously described method (42). Briefly, the last adenosine monophosphate of the 3’ end of the tRNA^Gly^ was exchanged at 25°C using alfa ^32^P-ATP (Amersham biosciences) and the CCA nucleotidyltransferase from *E. coli* (Eco-CCA) in a reaction containing 1 μM tRNA^Gly^_GCC_, 20 mM Tris-HCl pH 7.4, 40 mM NaCl, 18 mM DTT, 2.5 mM Na-pyrophosphate, 15 μM Eco-CCA, 5 μL of α^32^P-ATP. The reaction was allowed to proceed for 1 min, after which, Na-pyrophosphatase (Sigma Aldrich Cat. No. 322431) was added, and the reaction was allowed to continue for 4 more minutes. The reaction was stopped by mixing with acid phenol-chloroform, centrifuged, and the top layer was re-extracted. Finally, the radioactively labeled tRNA^Gly^_GCC_ was purified by gel extraction, concentrated, and kept at -20°C until needed.

### AtGlyRS Aminoacylation Assays

The activity of the recombinant At-GlyRS enzyme was assayed at 45°C in 40 μL reactions mixes containing 50 mM Tris pH 8.0, 5 mM DTT, 100 mM NaCl, 20 mM MgCl_2_, 5 mM glycine, 5 mM ATP, 1 μL of labeled tRNA^Gly^_GCC_ (∼2000CPM/μL), 2 μM tRNA^Gly^_GCC_, and 2 μM At-GlyRS. The addition of enzyme started reactions and time points of 1 μL were taken every 30 seconds, quenched using 5 μL of 0.4 M Na-acetate solution at pH 5.2 and 0.01 mg/mL P1 nuclease (SIGMA #N8630) to digest the aminoacylated tRNA^Gly^_GCC_. Once all the time points were taken, 10 minutes were allowed for the P1 nuclease to complete digestion. Then, 1 μL of the digested mix was spotted onto PEI-cellulose thin layer chromatography (TLC) plates (SIGMA #Z122882) and developed in a TLC chamber using 100 mM ammonium acetate and 5% acetic acid. Once the buffer reached the top of the TLC plate, they were air-dried and exposed to a phosphorimager screen inside a cassette for 16 h. The phosphor screen was scanned in a Typhoon Imager (Amersham), and the intensities of each spot obtained with ImageQuant 5.2 to plot the relation between reactants and products (Figure S14A). Under these conditions, reactions reached a plateau at 90 seconds, and the maximum aminoacylated fraction of tRNA^Gly^_GCC_ was 0.60 (Figure S14B).

### AtGlyRS-GSI Inhibition Assays

To verify the GSI’s ability to inhibit the aminoacylation reaction of At-GlyRS, the reactions were carried out under the same conditions as the aminoacylation reactions described above, except that 1 mM or 10 mM GSI were added to the reaction mix. Aliquots for points were also quenched, digested, and spotted onto PEI-cellulose plates and processed as the ones for the aminoacylation reactions (Figure S15).

### T7 RNA Pol system for Td-tRNA^Gly^ overexpression

We designed an *in vivo* tRNA overexpression system based on RNA Bacteriophage Polymerase T7 (T7 RNA Pol) to produce the substrate needed for our experiments. The methodology depends on overexpression being performed on a host producing T7 RNA Pol endogenously. Subsequently, this polymerase is responsible for the transcription of a sequence of DNA codifying for a tRNA gene located in an expression plasmid. Two different plasmids were designed and tested; the two differ in the tRNA sequence they encode (Figure S16). The tRNA sequences of *Escherichia coli* and *Thermanaerothrix daxensis* corresponding to the GCC anticodon were chosen. These plasmids were synthesized by the company ATUM. Once the product is transcribed, the host tRNA maturation machinery makes the necessary modifications to process the primary transcript (Figure S16).

BL21(DE3) (New England BioLabs, C2527I) cells transformed with the above plasmids were selected and cultured in Terrific Broth (TB) medium. Overexpression of the T7 RNA Pol was induced at an OD_600nm_ =0.6 with 1 mM IPTG. Cell pellets were collected by centrifugation to an RCF of 7480 × g after 24 hours of expression, then extracted with phenol, thus recovering the soluble RNA. The total soluble RNA extract is then subjected to liquid ion-exchange chromatography in an FPLC system at pH 5.0 in source Q medium using a buffer with Piperazine 20 mM and EDTA 250 μM. The tRNA was eluted with a gradient S-39 of 0 to 1 M NaCl, and the fraction of interest collected between 370 mM and 470 mM NaCl. The recovered fraction was applied to liquid chromatography of hydrophobic interaction with Toyopearl Butyl 650, equilibrated at pH 4.5 in 3.5 M ammonium sulfate, 50 mM sodium acetate, 5 mM MgCl_2,_ and 0.5 mM EDTA. The tRNA elutes at a staggered gradient of 3.5 M to 1.75 M of ammonium sulfate. The final fractions were exchanged in a storage buffer 50 mM Tris pH 7.5, 50 mM NaCl, 10 mM MgCl_2_, and concentrated at 60 mg/mL by ultrafiltration. Afterward, the sample was applied and stored at -20°C.

### Purification of Td-tRNA^Gly^ by probe hybridization

In general, a protocol previously reported in the literature (43) was followed with some modifications. Ultrafree-MC ultrafiltration tubes (0.22μm, Millipore UFC30GV0S) were loaded with 600 μL of Streptavidin Sepharose HighPerformance resin (GE Healthcare). To equilibrate the resin, 300 μL of a 10 mM Tris-HCl pH 7.5 solution treated with DEPC were added and mixed with a micropipette. Then, the excess solution was removed by centrifugation at an RCF of 18,000 × g for 10 seconds. This process was repeated three times to ensure that the resin was equilibrated at the desired pH.

The equilibrated resin was mixed with 400 μL of 10 μM 3’ biotinylated DNA oligo complementary to the target tRNA sequence. This solution was mixed with a micropipette and incubated for 10 minutes at room temperature. After incubation, the supernatant was removed by centrifugation at RCF of 18,000 × g for 10 s, and the resin was washed twice with 10 mM Tris-HCl pH 7.5. After removing the residual supernatant from the wash, the resin was resuspended in the total RNA fractions. The RNA was in hybridization buffer, 20 mM Tris-HCl (pH 7.5), 1.8 M NaCl, 0.2 mM EDTA. This mixture was incubated at 65°C for 10 min to denature the RNA and hybridize the DNA oligo. After incubation, the resin was allowed to equilibrate to room temperature.

To remove the RNA not attached to the stationary phase, the resin was washed by resuspension and centrifugation at 4°C with 400 μL of the 10 mM solution of Tris-HCl pH 7.5 until the recovered supernatant showed absorbance at 280 nm < 0.01 miliunits. The RNA was eluted by heating the resin, previously mixed with 300 μL of the desired storage buffer at 60°C for 5 min. Subsequently, the RNA was recovered by centrifugation at an RCF of 18,000 × g for 10 s.

### β-TdGlyRS-tRNA affinity measured by Biolayer Interferometry

The biolayer interferometry experiments described here were conducted on an eight-channel Octet RED96 equipment (FortandBio). Generally, we followed a standard protocol for this methodology (Tobias, 2013). In all experiments, Ni-NTA biosensors (NTA) (Fort and Bio) were used, thus taking advantage of the polyhistidine label of the protein. The concentration used to immobilize the β-bacGlyRS to the biosensor was 50 μg/mL. No temperature control was used. The steps and parameters used in all the experiments are described in the table below. It should be noted that the baselines and the dissociation step were made in different wells.

| Steps and parameters used in biolayer interferometry experiments. |  |  |  |
| --- | --- | --- | --- |
| Step | Procedure | Time(s) | Stirring (rpm) |
| 1 | Baseline | 60 | 1000 |
| 2 | Ligand load | 250 | 1000 |
| 3 | Baseline (2) | 800 | 1000 |
| 4 | Partnership phase | 200 | 1000 |
| 5 | Dissociation phase | 200 | 1000 |

To find the conditions that maximized the association of tRNA to β-bacGlyRS, a simple buffered solution -HEPES pH 7.5- was used. Different magnesium salts - acetate, nitrate, chloride, sulfate, and magnesium nitrate - were tested in a range of concentrations between 0 and 200 mM. We also experimented with the effect of several monovalent salts - chlorides and nitrates of sodium and potassium, as well as ammonium acetate. Spermine, spermidine, and glycine were tested in a range of concentrations between 5 and 50 mM. The optimized binding kinetics experiment was finally performed in 50 mM HEPES pH 7.5, 50mM MgCl_2_, 5 mM ammonium acetate, and 5 mM spermine. The tRNA concentrations tested were 0, 0.5, 1, 3, 10, 25, 50, and 100 μM. Data analysis was done in the Octet ^®^ Data Analysis HT (ForteBio) program.

### Single-particle electron microscopy by negative staining

#### - Preparation of biological samples

Prior to grid preparation, the apo sample was dialyzed overnight at room temperature against buffer containing 50 mM HEPES pH 7.5, 200 mM NaCl and 10 mM MgCl_2_. The dialyzed sample was then diluted to a final concentration of 15 ng/µL.

Commercially available carbon coated grids (Electron Microscopy Sciences, CF200-Cu) were cleaned utilizing a plasma cleaning system (Gatan Solarus 950) operating at 25 W for 10 seconds. Apo bacGlyRS (15 ng/ µL) was loaded by allowing the grid to float carbon-layer side down on a 30 µL droplet of sample for 5 minutes. Subsequently, washing was accomplished by transferring the grid to a 30 µL droplet of dialysis buffer for 1 minute; this process was performed twice. After washing, the grid was dried by side blotting and transferred to a 1% uranyl formate droplet to be incubated for 30 seconds. Side blotting and incubation on uranyl formate was repeated once more afterwards. Lastly, the grid was side blotted again and allowed to dry at room temperature for 3 minutes.

#### - Data collection and processing

Automated data acquisition was performed at a 0° tilt angle with 30,000x magnification at an average defocus of -0.9 μm on a JEOL 1400 microscope at 120 kV with 4k x 4k CCD (Gatan Ultrascan), using the program Leginon (21). Data processing was attained by following the general protocol detailed in the RELION 3.1 software tutorial (44). Briefly, ∼1,500 particles were selected manually to get templates for autopicking. The particles were then subjected to reference-free alignment followed by 2D classification into 7 classes (Figure S21). De novo 3D model generation from the 2D particles was achieved using RELION’s Stochastic Gradient Descent (SGD) algorithm, followed by unsupervised 3D classification. Refined volumes were estimated to be in the resolution range of 21.8-25.0 Å using the Fourier Shell Correlation (FSC) gold standard. The reconstructed envelope was fitted to the tetramer model obtained by X-ray crystallography by manually placing them together and subjecting them to automated optimization in the program CHIMERA (Figure S22) (45). Some parameters of the collected data are shown in Table S5.

